# Molecular studies into Copine-4 function in Retinal Ganglion Cells

**DOI:** 10.1101/2021.08.09.455730

**Authors:** Manvi Goel, Angel M. Aponte, Graeme Wistow, Tudor C. Badea

## Abstract

The molecular mechanisms underlying morphological diversity in retinal cell types are poorly understood. We have previously reported that several members of the Copine family of Ca-dependent membrane adaptors are expressed in Retinal Ganglion Cells (RGCs) and transcriptionally regulated by Brn3 transcription factors. Several Copines are enriched in the retina and their over-expression leads to morphological changes reminiscent of neurite formation in HEK293 cells. However, the role of Copines in the retina is largely unknown. Here we focus on Cpne4, a Copine whose expression is restricted to RGCs. Over-expression of Cpne4 in RGCs in vivo led to formation of large varicosities on the dendrites but did not otherwise visibly affect dendrite or axon formation. Protein interactions studies using yeast two hybrid analysis from whole retina cDNA revealed two Cpne4 interacting proteins - HCFC1 and Morn2. Mass Spectrometry analysis of retina lysate pulled down using Cpne4 or its vonWillebrand A (vWA) domain identified a further 207 interacting proteins. Gene Ontology (GO) analysis of Cpne4 interactors suggests its involvement in several metabolic and signaling pathways, including processes related to vesicle trafficking, intracellular membrane bound organelles, and plasma membrane associated structures, including neurites. We conclude that, consistent with its domain structure, Cpne4 may be involved in assembly and trafficking of several membrane associated cell compartments.

## 1. INTRODUCTION

RGCs in the retina transmit visual signals received by photoreceptors to the brain for processing visual inputs. Different RGC sub-types are responsible for computing different aspects of the visual stimuli. The combinatorial expression of transcription factors in different RGC sub-types regulates cell specific morphologies and physiology by controlling molecules involved in the development of dendrite/axon morphology, synapse formation and function (1,2).

How cell-specific morphologies develop in the retina is not well understood. One likely mechanism is that transcription factors encode specific morphological features via adhesion molecules or cytoskeletal elements they regulate. For example, Tbr1 regulates cell adhesion molecules Cdh8 and Sorcs3 resulting in dendritic stratification of J-RGCs in the Off sublamina (3). Other such molecules might be responsible for cell specific morphologies in other RGC types. Copines are a family of such candidate cell morphology determinants, regulated by Brn3 transcription factors. Copines consist of two C2 domains (C2A and C2B) and a vWA domain (4,5). The Copine C2 domains are similar to those found in a variety of vesicular traffic proteins such as Synaptotagmins, Munc18, Rabphilin3A and Doc2, and have been involved in calcium dependent binding to cell membranes (6). vWA domains are typically found extracellularly in several proteins (e.g. integrins) and are involved in protein-protein interactions (7). However Copines (and their vWA domain) are intracellular proteins that interact transiently with the inner leaflet of the plasma membrane.

There are nine Copines in mammals-Cpne1-9 (4,8–10). Of these, Cpne1, 2 and 3 are expressed ubiquitously. Cpne4, 5, 6, 7, 8 and 9 are enriched in neurons. Cpne5 and 8 are also expressed in other tissues such as kidneys, lungs, testes, mammary glands (11,12).

Copines are conserved across several species. They have previously been shown to be important in a variety of cellular functions. Copines are involved in myofilament stability in C. elegans and plant growth in Arabidopsis (13,14). Copines are also involved in cytokinesis and contractile vacuole function by regulating cAMP signaling in Dictyostelium (15). Copine has also been shown to interact with actin filaments to regulate chemotaxis and adhesion in Dictyostelium (16).

In the nervous system, Copines were first seen to be localized in hippocampus and olfactory bulb neurons in mouse brain (10,17). Cpne6 is expressed in the hippocampus and is required for regulating the spine morphology during long term potentiation in hippocampus by regulating the Rac-LIMK-Cofilin and BDNF-TrkB pathways (18,19). Cpne1 has been previously seen to be upregulated during development and is required for hippocampal progenitor cell differentiation into neurons (20,21). Cpne7 is expressed in sublaterodorsal nucleus in pontine segmental area and is required for REM sleep (22).

In the retina, we have previously reported that Cpne4, 5, 6 and 9 are enriched in the inner retina and they are regulated by Brn3b and Brn3a (1,23). Whereas Cpne5, 6 and 9 were expressed in most of the GCL as well as INL, Cpne4 is the only Copine specifically expressed in RGCs (with the exception of one INL amacrine cell type) (23). Using over-expression studies in HEK293 cells, we found that Copines can significantly alter cell morphology, inducing elongated membrane processes similar to neurites. In the current study, we further explore the effects of Cpne4 expression in RGCs and study its protein interactions using yeast two hybrid analysis and Mass Spectrometry, to identify its cell biological functions and role in RGCs.

## 2. MATERIALS AND METHODS

### 2.1 Transfection in HEK293-Cre cells

The cDNA for full length Cpne4, two C2 domain or vWA domain were cloned into pAAV-FLEX-HA-T2A-meGFP plasmid vector (Fig.1A). The cDNA is in frame with 3XHA (3 tandem copies of Hemagglutinin antigen tag-5’ TACCCATACGATGTTCCAGATTACGCT 3’ or 5’ TATCCATATGATGTTCCAGATTATGCT 3’) and separated from meGFP (membrane eGFP) by a P2A peptide sequence. The constructs were transfected into HEK293-Cre cells (a kind gift of Dr. Brian Sauer) [1, 23, 62] using Lipofectamine (Invitrogen, Carlsbad, CA). The transfected cells were fixed in 2%PFA after 48 hours and processed for immunofluoroscence as described in section 2.4.

**Figure 1:**
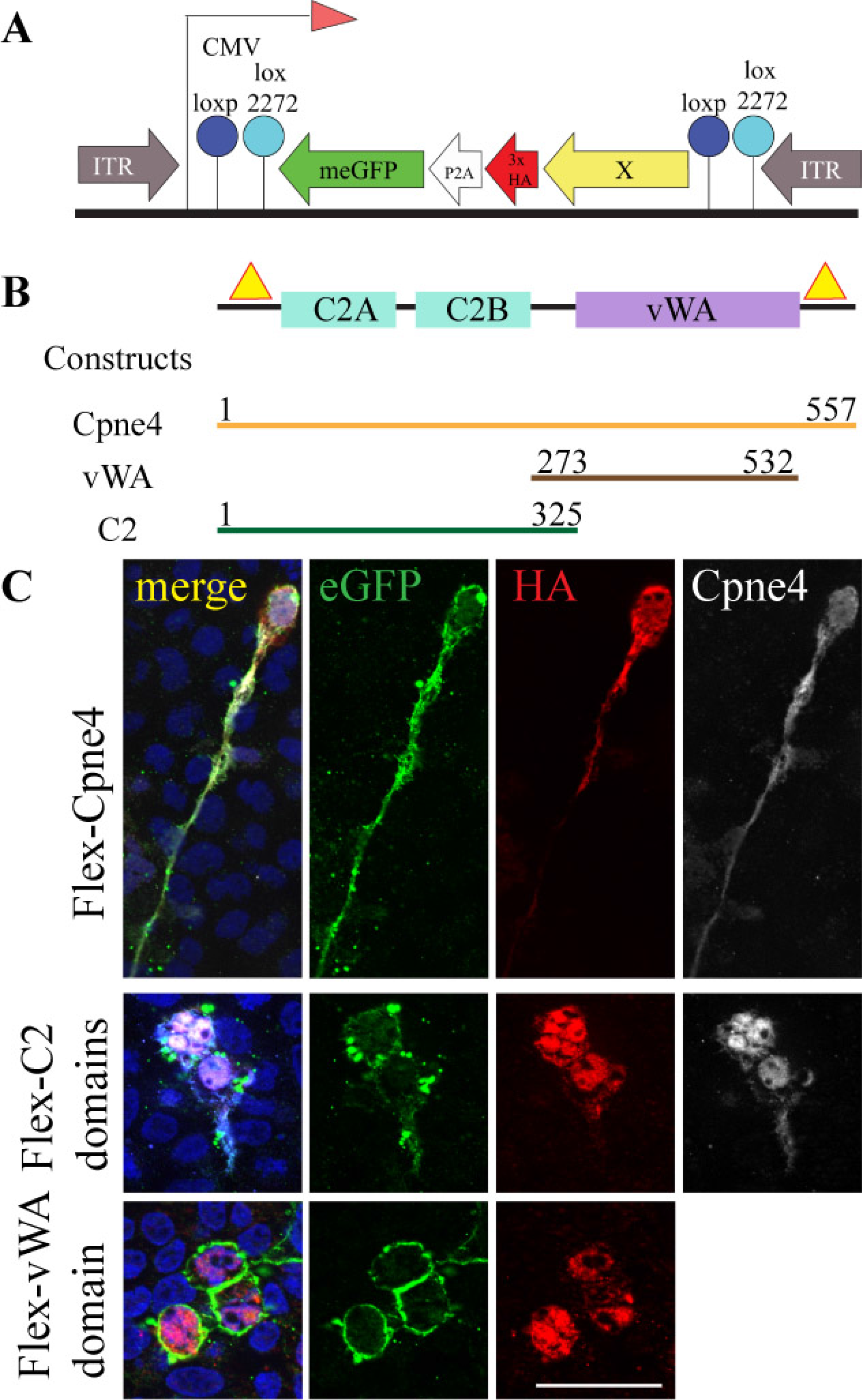
Cpne4 and Cpne4 dominant negative transfections in HEK293: (A) Map of the Flex construct. (B) Map of domain structure of Cpne4 shows three domains of the Cpne4 protein-two C2 domains (blue) and a vWA domain (purple). The location of the three Cpne4 plasmid constructs are shown by full length Cpne4 construct (orange), vWA domain construct (brown) and C2 domains construct (green). The binding of the two Cpne4 antibodies-N-terminal and C-terminal antibodies on the Cpne4 protein are indicated by yellow triangles. (C) Representative images of HEK293 cells transfected with expression constructs for full length Cpne4 (top row), C2 domains construct (middle row) and vWA domain construct (bottom row). The cells were counterstained for eGFP (green), HA (red) and N-terminal or C-terminal Cpne4 (white) antibodies. Scale bar: 50um.

### 2.2 Mouse lines

Adult *Brn3b^KO/KO^* (or KO) (24) and *Brn3b^W/WT^* (or WT) littermates were used for immunohistochemistry (IHC). Postnatal day 0 (P0) or P14 *Brn3b^Cre/WT^* mice were used for AAV1 virus infections. Adult wild-type (C57/Bl6 – SV129 mixed background) mice were used for retina pulldowns and mass spectrometry experiments, to determine the protein interactors of Cpne4. All animal procedures were approved by the National Eye Institute (NEI) Animal Care and Use Committee under protocol NEI640.

### 2.3 Virus infections in retina

Flex-meGFP-P2A-HA-Cpne4, described in section 2.1, was packaged into AAV1 viruses (henceforth AAV1-Cpne4). Intraocular injections of 0.5 ul (1e9 viral particles/ul) of AAV1-Cpne4 or AAV1-eGFP control virus were done in P0 or P14 Brn3bCre/WT mice eyes using glass capillaries fitted onto a Femtojet device (Eppendorf, Enfield, CT), as previously described. Injections were aimed at the scleral region adjacent to the limbus. Eyes were collected at 2-3 months of age and flat-mounted for immunostaining. The eyes were fixed in 4% paraformaldehyde for 15 minutes and retinas were dissected to make a flat mount preparation. The retinas were again fixed for 30 minutes and then washed three times with PBS+ 0.5% Triton-X 100 (PBST). Immunofluorescence was performed as described below.

### 2.4 Immunofluorescence

Brn3b WT and KO sections were co-immunostained to confirm the presence of different interactor proteins identified from yeast two hybrid and mass spectrometry analysis in the retinal ganglion cells. Similar process was followed for immunofluorescence of transfected HEK293 cells on coverslips. The sections or cells were incubated with blocking solution-10% bovine serum albumin (BSA), 10% normal donkey serum (NDS) and 0.5% Triton X 100, for one hour at room temperature. The blocking solution was then replaced with primary antibody solution containing the antibodies at the required concentrations and incubated overnight, at 4^0^C. Sections or cells were washed three times with PBST and incubated with the secondary antibody solutions for one hour at room temperature. The sections or cells were washed again with PBST and coverslipped.

For staining the AAV1 infected retina flatmounts, the retinas were incubated in blocking solution, overnight at 4^0^C. The blocking solution was then replaced with primary antibody solution and retinas incubated for 48 hours at 4^0^C. The retinas were then washed three times in PBST, secondary antibody solution was added and retinas were incubated overnight at 4^0^C. The retinas were washed again three times with PBST, carefully mounted on glass slides and coverslips were placed. Image acquisition was on either a Axioimager Z2 fitted with an apotome device, or on a LSM 880 confocal microscope (both from Zeiss, White Plains, NY). The images were taken as 1um thick z-stacks and the images were stacked using ImageJ. Colocalization analysis of Copine4-interacting proteins in HEK293 cells was performed using the “Coloc 2” plugin in ImageJ.

### 2.5 Yeast two hybrid

A Gal4 based yeast two hybrid analysis was performed to identify the proteins that interact with Cpne4 vWA domain. An adult mouse retina cDNA library was cloned in pGADT7 (carrying a Trp1 selection gene) and transformed in AH109 yeast strain (containing His3, Ade2, lacZ and Mel1 selections; Clontech, BD Biosciences, Pao Alto, CA). Cpne4 vWA domain was cloned into pGBKT7 (carrying Leu2 selection gene) and transformed in to competent Y187 yeast strain (lacZ and Mel1 selection). The yeast mating experiment was performed as per the Two-hybrid library screening protocol for yeast mating (Clontech). Briefly, one colony of Y187 transformed with pGBKT7+ Cpne4 vWAdomain was inoculated in 50 ml of Tryptophan (Trp) selection media and incubated at 30^0^C overnight. The following day, the culture media was centrifuged, and the pellet re-suspended in 5ml Trp selection media. 45ml of YPDA media (YPD media supplemented with adenine(Ade)) was added to it. 1ml of cDNA library (titer= 5 x 10^7^cfu/ml) was thawed in a water bath at room temperature and added to the above and allowed to mate for approximately 24 hours at 30^0^C with slow shaking at 50rpm. The next day, the presence of mated, diploid yeast cells was checked under light microscope. The culture was then centrifuged, and the pellet resuspended in 15ml 0.5X YPDA. 300ul aliquots of the entire 15ml mating culture was spread on 15cm quadruple selection agar plates (with selection for Trp, leucine (Leu), Ade and histidine (His) and X-alpha-Gal reporter) and grown for about 5 days at 30^0^C. Small scale positive and negative control matings were also performed. For the positive control, pGBKT7-53 encoding the p53 protein and pGADT7-T encoding the SV40 large T antigen protein were transformed into Y187 and AH109, respectively. For negative control, empty pGBKT7 and pGADT7, with no gene insertions, were transformed into Y187 and AH109, respectively. For the positive control mating, one colony each from Y187 + pGBKT7-53 and AH109 + pGADT7-T were inoculated in 500ul 2X YPDA and incubated overnight at 30 degrees at 200rpm. Similarly, for the negative control mating, Y187 + empty pGBKT7 and AH109 + pGADT7 were inoculated in 500ul 2X YPDA and incubated overnight at 30 degrees at 200rpm. The following day the control cultures were spread in 1:10, 1:100 and 1:1000 dilutions on separate single (Leu or Trp), double (Leu and Trp) or quadruple selection agar plates and grown for 3-5 days at 30^0^C.

After 4 days of selection, 241 pale blue colonies were picked for confirmation. They were streaked separately on fresh agar plates with quadruple selection. Six colonies grew into blue colonies after 3-4 days. For both positive and negative controls, the colonies appeared on the single selection and double selection plates. But on the quadruple selection plates, blue colonies appeared only on the positive control and there were no colonies for the negative control. Colony PCR was performed on the six selected colonies, and PCR products were extracted from the gel and Sanger sequenced (Eurofins, Luxembourg). Inserts were identified by BLAST searches against the NCBI mouse transcriptome collection.

### 2.6 Co-immunoprecipitation

HEK293-Cre cells were co-transfected with HA-eGFP-Cpne4-vWAdomain construct (described in section 2.1) and Flag-tagged target protein or protein domain identified from yeast two hybrid analysis. 24 hours after the transfection, the cells were washed three times with 1X PBS. 300ul lysis buffer (50mM Tris-HCl, 150mM NaCl, 0.5% NP40 and 1mM EDTA) containing protease inhibitor (Roche, Basel, Switzerland) and 0.2M PMSF was added to each well and incubated on ice for 15 minutes. The lysate was then collected in 1.5ml tubes and centrifuged at 14000g for 10 minutes to remove any debris. Meanwhile, magnetic beads were washed with 1X PBS and incubated with Flag antibody for 30 minutes, with end-to-end rotation at 4^0^C. The beads were again washed with 1X PBS to remove any unbound antibody and the supernatant from the cell lysate was added to it. These were incubated overnight with end-to-end rotation at 4^0^C. The next day, the beads were washed four times and PBS + Laemmli buffer (62.5mM Tris-HCl, 2% SDS, 10% glycerol, 5% beta-mercapto-ethanol, bromophenol blue) was added to the beads. The beads were boiled for 5 minutes at 70^0^C. The beads were separated on a magnetic rack and the supernatant loaded on a 10% SDS-PAGE gel. The gel was allowed to run until the dye front reached the bottom of the gel. The separated proteins were then transferred to 0.2um PVDF membranes and processed further for Western blotting.

### 2.7 GST pulldown from retina

GST tagged Cpne4 protein was synthesized from bacteria as described before (23). GST-Cpne4-vWAdomain and GST proteins were also synthesized similarly. 19 wild-type retinas (C57Bl6 and SV129) were homogenized using a glass homogenizer, in cold lysis buffer (RIPA buffer: 50mM Tris-HCl, 150mM NaCl, 1mM EDTA, Complete protease inhibitor (Millipore Sigma, Burlington, MA; Catalog no. 11697498001)). NP40 was then added to the lysate to a 0.5% final concentration. Glutathione-tagged magnetic beads were added to the lysate and incubated at 4 degrees with end-to-end rotation for 2 hours. The magnetic beads were removed, and the lysate centrifuged at 700g for 10 minutes to remove any debris. The lysate was then kept on ice until further process. Equimolar amounts of GST, GST-vWA domain and GST-Cpne4 were incubated with glutathione tagged magnetic beads and incubated with end-to-end rotation at 4^0^C for 2 hours. The supernatant was discarded, and beads washed three times with PBS. Equal volume of cleared retina lysate (600ul each) was added to each of the above three tubes and incubated overnight at 4^0^C with end-to-end rotation. The following day, the supernatant was discarded, and the beads were washed four times with PBS. 30ul of glutathione elution buffer was added to each of the tubes and incubated at room temperature for 10 minutes with end-to-end rotation. Laemmli buffer was added and the samples boiled at 70^0^C for 5 minutes. The beads were separated, and the samples loaded onto 4-15% gradient SDS-PAGE gels. The samples were allowed to run on the gel until the dye front reached the bottom and proteins were visualized using Coomassie blue staining. Full lanes were cut out of the gel for each of the three samples, cut into smaller band size pieces and collected in separate tubes. Three such replicates were prepared each for GST, GST-Cpne4-vWAdomain and GST-Cpne4 pulldowns. The cut bands were then processed further for liquid chromatography mass spectrometry (LC-MS). An additional three replicates also prepared similarly, transferred to PVDF membranes directly after SDS-PAGE and checked with specific antibodies by western blotting. See the section on western blotting for further details.

### 2.8 Mass spectrometry

For sample preparation for LC-MS, Coomassie blue stained protein gel bands were first destained with 30% ethanol solution until the gel pieces became transparent. The gel pieces were then destained in 65% methanol + 10% acetic acid solution for 30 minutes. The destaining solution was removed and bands were washed with 100 mM TEAB (tri-ethyl ammonium bicarbonate) solution for 5 minutes. 250 ul dehydration solution (75% acetonitrile in 100 mM TEAB) was added and the bands were agitated for 10 minutes at room temperature. The dehydration solution was removed, and the bands were air-dryed for a couple of minutes. The gel pieces were then dehydrated with reducing solution (10 mM TCEP in 100 mM TEAB) for 45 minutes at 56^0^C. This solution was replaced by dehydration solution and bands incubated for 10 minutes at room temperature. The bands were then air dried and alkylation solution (20 mM iodoacetamide in 100 mM TEAB) was added and incubated for 30 minutes at room temperature. The gel bands were washed again and dehydrated again in the dehydration solution until the gel bands shrunk to half the size. The gel bands were air dried briefly and trypsin solution was added to the bands, enough to cover the bands. The bands were left on ice for 5 minutes and more trypsin was added as needed. After 5 minutes, the trypsin solution was removed and 100 mM TEAB was added to tubes, enough to cover the bands. The tubes were incubated at 37^0^C for overnight digestion. The next day, the trypsin solution was removed, and digested peptides transferred to new tubes separately for each sample. 150ul of extraction solution (75% acetonitrile, 0.1% formic acid) was then added to each of the tubes and agitated for 10 minutes at room temperature. A short spin was done at 10,000 g and the gel pieces were saved. The extracts were then vacuum dried and 25 ul 1% trifluoroacetic acid was added to each tube. This was followed by a peptide clean-up as per the zip-tip protocol (Millipore Sigma; Cat# C18 ZTC18S008). 20ul of 0.1% formic acid (in acetonitrile) was aspirated in the zip-tip and discarded. 20ul of 0.1% formic acid (in water) was similarly aspirated and discarded. The peptide extract was then aspirated 7-10 times, to let the peptides bind to the zip-tip column. This was followed by washing the zip-tip three times by aspiring 20ul of wash solution (0.1% formic acid in water) and dispensing it. To elute the bound peptides, 20ul of elution buffer (75% acetonitrile + 0.1% formic acid) was aspirated and dispensed in a tube. This step was repeated five times and each time eluate was collected in the same tube. The tubes were then dried in a speed vacuum. 20ul of 2% acetonitrile + 0.1% formic acid solution was added to the dried peptide digest and the tubes were vortexed and centrifuged. The solution was then transferred to LC vials for LC-MS analysis.

Desalted tryptic peptides were analyzed using nanoscale liquid chromatography tandem mass spectrometry (nLC-MS/MS) and Ultimate 3000-nLC online coupled with an Orbitrap Lumos Tribrid mass spectrometer (Thermo Fisher Scientific, Waltham, MA). Peptides were separated on an EASY-Spray Column (Thermo Fisher Scientific; 75 μm by 50 cm inner diameter, 2-μm particle size, and 100-Å pore size). Separation was achieved by 4 to 35% linear gradient of acetonitrile + 0.1% formic acid for 90 minutes. An electrospray voltage of 1.9 kV was applied to the eluent via the EASY-Spray column electrode. The Orbitrap Lumos was operated in positive ion data-dependent mode. Full-scan MS was performed in the Orbitrap with a normal precursor mass range of 380 to 1500 *m/z* (mass/charge ratio) at a resolution of 120,000. The automatic gain control (AGC) target and maximum accumulation time settings were set to 4 × 10^5^ and 50 ms, respectively. MS was triggered by selecting the most intense precursor ions above an intensity threshold of 5 × 10^3^ for collision-induced dissociation (CID)–MS fragmentation with an AGC target and maximum accumulation time settings of 5 × 10^3^ and 300 ms, respectively. Mass filtering was performed by the quadrupole with 1.6 m/z transmission window, followed by CID fragmentation in the ion trap (rapid mode) and collision energy of 35%. To improve the spectral acquisition rate, parallelizable time was activated. The number of MS spectra acquired between full scans was restricted to a duty cycle of 3s.

Raw data files were processed with the Proteome Discoverer software (v2.4, Thermo Fisher Scientific), using Sequest HT (Thermo Fisher Scientific) search node for carbamylated peptide/protein identifications. The following search parameters were set: protein database UniProtKB/Swiss-Prot *Mus musculus* (17,033 sequences release 2020_10) concatenated with reversed copies of all sequences; MS1 tolerance of 12 ppm: ion trap detected MS/MS mass tolerance of 0.5Da; enzyme specificity set as trypsin with maximum two missed cleavages; minimum peptide length of 6 amino acids; fixed modification of Cys (carbamidomethylation); variable modification of methionine oxidation and acetyl on N terminus of protein. Percolator algorithm (v.3.02.1, University of Washington) was used to calculate the false discovery rate (FDR) of peptide spectrum matches (PSM), set to a q-value <0.05(2–5)

In order to identify retinal proteins that bind differentially to full-length Copine4 or Copine4-vWA domain, we compared the peptides pulled down by either GST-vWA, GST-Copine4 or GST control identified in the previous step. Of the 2119 proteins pulled down in either of the three conditions, we selected for further analysis those that were represented in all three replicates of at least one condition. Differential display analysis was performed using the R package “DEP” (DEP 1.12.0, <u>10.18129/B9.bioc.DEP</u>; (25)). Missing values were replaced with 0, and data was normalized using VSN normalization. Pairwise comparisons for GST only vs. GST-Cpne4 and GST-vWA samples was performed at thresholds of 0.05 FDR and 2-fold change.

### 2.9 Western blotting

Western blotting (WB) was done as described before (23). Briefly, the PVDF membranes were washed with Tris buffered saline with 0.1% Tween20 (TBST). The membranes were then incubated in 5% milk (in TBST) for 1 hour at room temperature. Primary antibody solution prepared in 5% milk was added to the respective membranes and incubated overnight at 4C on a rocker shaker. The next day, the membranes were washed three times in TBST and secondary antibody solution (in 5% milk) was added. The membranes were then incubated at room temperature for 1 hour, followed by three washes with TBST. The membranes were exposed to Super signal chemiluminescence (Thermo Fisher Scientific) for 5 minutes and images taken on a gel dock (Bio-Rad, Hercules, CA).

### 2.10 Statistical analysis

The statistical analysis for comparing the varicosity areas in Cpne4 with control infected RGCs was done using Kolmogorov-Smirnov test (KS2, Matlab). For measuring colocalization of Cpne4 to interacting proteins in co-transfected HEK293 cells, Spearmann’s correlation coefficient was used. Number of images, animals and cells measured and p-values are given in the corresponding Results sections.

## 3. RESULTS

### 3.1 Cpne4 transfection induces morphology changes *in-vitro*

To evaluate the subcellular distribution of Copine4 or its domains, we transfected expression vectors containing full length Cpne4, C2 domains or vWA domain tagged with HA (Fig.1A, B), in conjunction with a plasma membrane attached eGFP variant (meGFP) into HEK293 cells. Full length Copine4, C2 domains or vWA domain were detected in the nuclei, cell body as well as plasma membrane of the cells (Fig.1C) as indicated by both HA (red) and Cpne4 (white) staining. As previously reported, full length Cpne4 induces morphological changes (Fig. 1C top panel; n=2 experiments) resembling neurites (23). C2 domain or vWA domain constructs also induce extended process formation occasionally, although morphological changes were not as prominent and wide spread as with Cpne4 transfection (Fig.1C middle and bottom panel; n=2 experiments each).

### 3.2 Cpne4 infection induces morphology changes *in-vivo* in RGCs

Given the morphological defects observed when expressing Copines in HEK293 cells in culture, we asked whether overexpressing Cpne4 in RGCs will affect RGC morphology. We therefore infected either P0 or P14 Brn3b^Cre/WT^ mice with AAV1 vectors co-expressing HA-tagged Cpne4 and meGFP in a Cre-dependent manner. Membranes from infected Brn3b^Cre/WT^ RGCs were labelled with eGFP (green), revealing axonal arbors, cell bodies and dendrites (Fig.2A, B). In contrast, Cpne4 expression as seen by HA (red) labeling was largely confined to the cell body and dendrites, and only rarely reached into the axon (Fig.2A, B; n=6 retinas, 72 cells). Interestingly, several large varicosities were observed on dendrites of Cpne4 infected Brn3b^Cre/WT^ RGCs. These large varicosities (we call these ‘blebs’, Fig. 2A, 2A1) can be easily distinguished from regular varicosities, as seen on the RGC dendrites in Cpne4 infected cells (Fig. 2A, 2A2, supplementary Figure 1A, n=6 animals, 72 cells), or on the dendrites of control RGCs infected solely with AAV1-eGFP (Fig.2C, E; area= 10.7± 0.61um^2^ for Cpne4 versus 3.12± 0.18um^2^ for controls, KS2 test p value < 0.05; controls n=8 animals, 73 cells). The size and number of blebs varied significantly across the infected RGCs, and bleb size tended to increase with RGC dendritic arbor area (Supplementary Figure 1B, C). A comparison between the P0 and P15 Cpne4 infected RGCs suggest no differences in the bleb areas (P0 mean area= 9.95± 1.11um^2^, P15 mean bleb area= 15± 3.75um^2^; p=0.2), suggesting that the process disrupted by Cpne4 overexpression is still active in nearly adult animals (P14), as opposed to the early stages of dendrite formation (P0). A high resolution (Zeiss LSM 880 airy scan) imaging revealed that blebs contained large amounts of Cpne4, as well as cell membranes, as revealed by membrane attached eGFP. Besides the formation of large varicosities or blebs, Cpne4 over-expression into RGCs did not cause any changes in the arbor area or stratification when compared to controls infected with eGFP (Fig.2F, G). Infection of RGCs with AAV1 vectors expressing either C2 domains or vWA domains alone did not cause significant morphological changes or formation of bigger blebs (data not shown).

**Figure 2:**
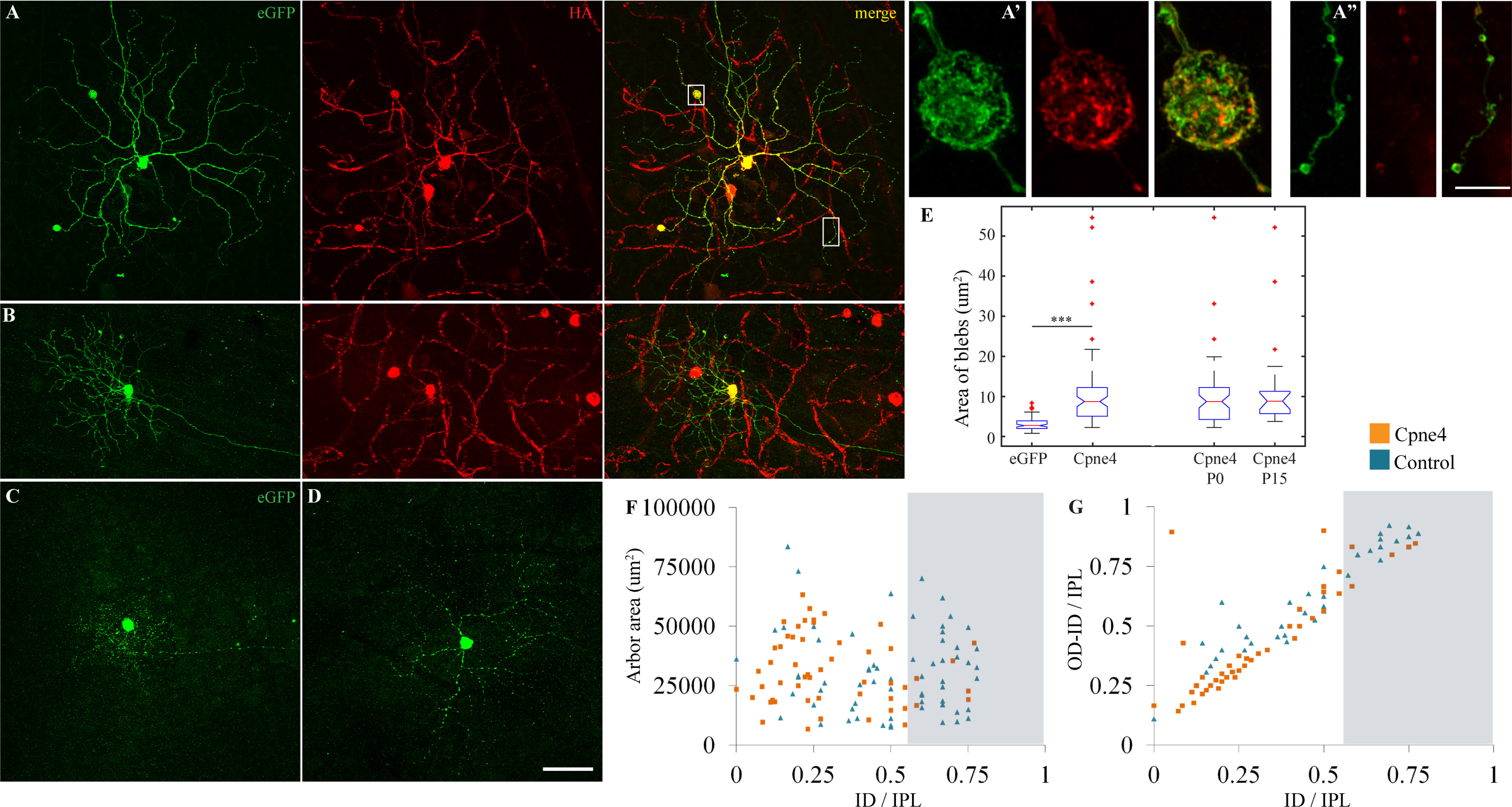
Cpne4 virus infections lead to ‘bleb’ formation on dendrites of Brn3b+ RGCs. (A, B) Intraocular injections of AAV1-Cpne4 viruses were done in P0 or P14 Brn3b^Cre/WT^ mice eyes. Blebs were seen occasionally on dendrites of Cpne4 infected RGCs. For controls, AAV1-eGFP was similarly injected in P0 Brn3b^Cre/WT^ mice (C, D). No blebs were observed in the control infected RGCs. Airy scan images for one of the blebs (A1) and adjoining dendrites (A2). (E) Measurement of the ‘bleb’ areas for Cpne4 infected RGCs as compared to regular varicosities on controls shows a significant difference between them while no difference between P0 and P15 Cpne4 infected retinas. (F, G) Areas and lamination measurements indicate no differences between the Cpne4 transfected RGCs as compared to control transfected RGCs. (ID: inner distance-distance between outermost tip of RGC dendrites and the INL; IPL: width of the IPL). Scale bar for A-C: 50um; Scale bar for A’, A”: 5um.

### 3.3 Yeast two hybrid analysis reveals Cpne4 vWA domain interaction with Morn2, HCFC1 and Tox3

The domain structure of Copines, and previous work on other family members suggests that Copine C2 domains facilitate Ca^2+^ mediated membrane attachment, while Copine vWA mediates interactions with other proteins. We therefore used a Gal4 based yeast two hybrid system using the Cpne4 vWA domain as bait protein and an adult mouse retina cDNA library as pray, in order to identify potential retina-specific interactors of Cpne4. Only five clones selected from the initial screening confirmed upon replating (Fig.3A). A colony PCR (Fig.3B) and subsequent Sanger sequencing on the PCR products identified the clones as domains of five proteins-2 clones of HCFC1, 1 each of Morn2, Tox3 and GAPDH and a mixed clone consisting of Rhodopsin and GAPDH (Fig.3B). Interestingly, the interacting domain of Tox3 protein contained the HMG box and that for HCFC1 contained an HCF repeat4. For Morn2 the interacting domain consisted of full length of the Morn2 protein and some non-coding sequence on the 5’ end (Fig.3C). Complete cDNA sequences of the regions of Morn2, HCFC1 and Tox3 interacting with Cpne4 are given in the Supplementary document.

**Figure 3:**
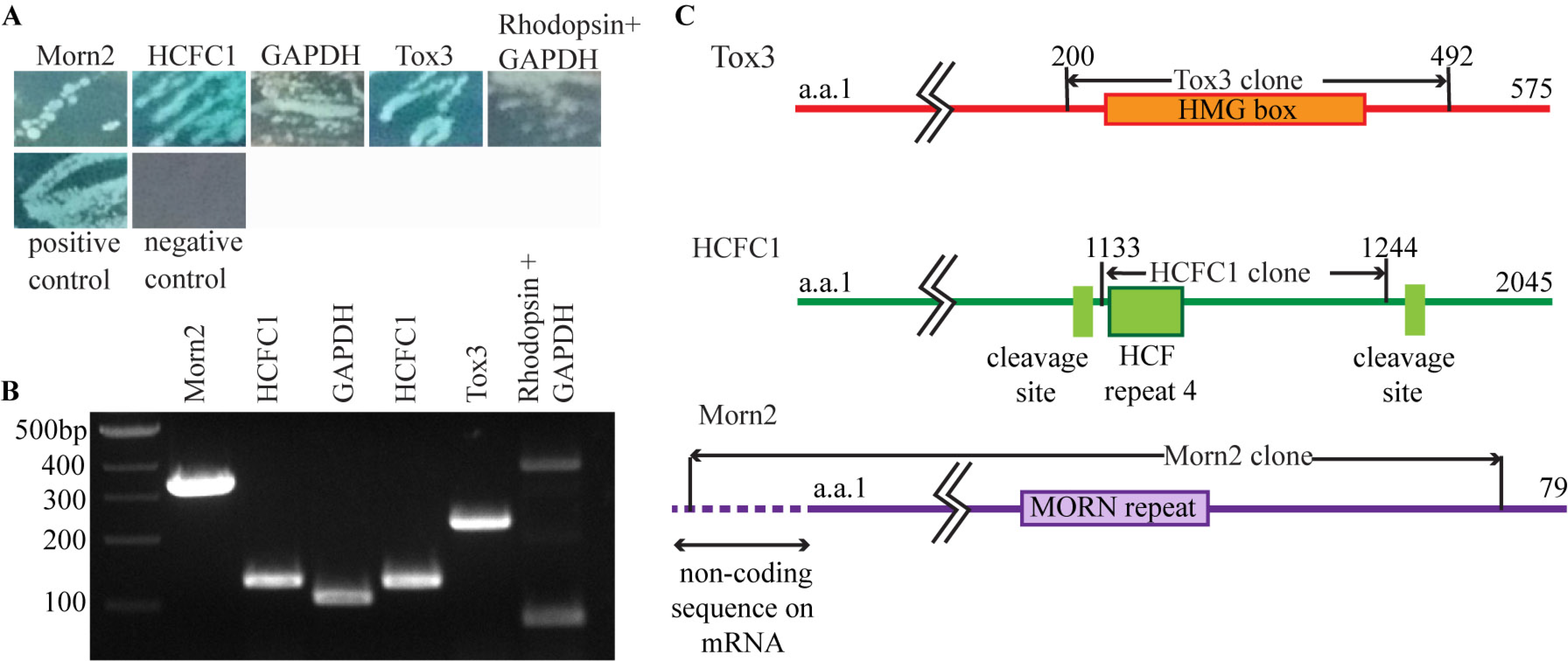
Yeast 2 hybrid analysis shows 5 proteins interacting with Cpne4 vWAdomain: (A) A Gal4 based yeast two hybrid analysis on the adult mouse retina cDNA library with Cpne4 vWA domain as bait protein showed five positive interactions (blue colonies). (B) Gel image of colony PCR performed on the the positive clones from the yeast two hybrid shows the corresponding interacting DNA sequences. The identity of these DNA sequences as confirmed by the Sanger sequencing followed by BLAST analysis are shown as labels on the lanes. (C) Sequences of some of the interacting peptides as identified by Sanger sequencing followed by BLAST analysis are shown-HCFC1 (green); Mycbp2 (orange) and Morn2 (purple). The relative positions of the actual interacting region on the respective protein is indicated by arrows and any protein domains in that region and surrounding regions are also labeled.

### 3.4 Cpne4 vWA domain interacts with HCFC1, Morn2 and Mycbp2 in HEK293 cells

We next sought to validate our candidate Cpne4 interactors (HCFC1, Morn2 and Tox3) by co-immunoprecipitation. In addition, we surveyed candidate Cpne4-vWA interactors identified in a previous yeast two hybrid analysis using a mouse embryonic cDNA library (26). By exploring our RNA sequencing analysis (1), we found that some of these previously reported candidate Cpne4 interactors (Bicd2, Pitpnm2, Sptbn1 and Mycbp2) were indeed expressed in RGCs (Supp. Fig. 2, Fig. 5A). We therefore cloned the interacting peptide regions from our candidates (HCFC1, Morn2 and Tox3) as well as the matches from the Tomsig et al. study (Bicd2, Pitpnm2, Sptbn1 and Mycbp2) downstream of the Flag tag in an eukaryotic expression vector and co-transfected them with the HA-tagged Cpne4-vWA in HEK293 cells.

The colocalization of these proteins was observed by immunostaining (Fig.4B, C, and E; n=6, 5 and 4 for Morn2, HCFC1 and Tox3 respectively). While Mycbp2 and Morn2 were localized in the cell body of the cells, Tox3 was always localized in the nuclei (Fig.4E; n=6).

**Figure 4:**
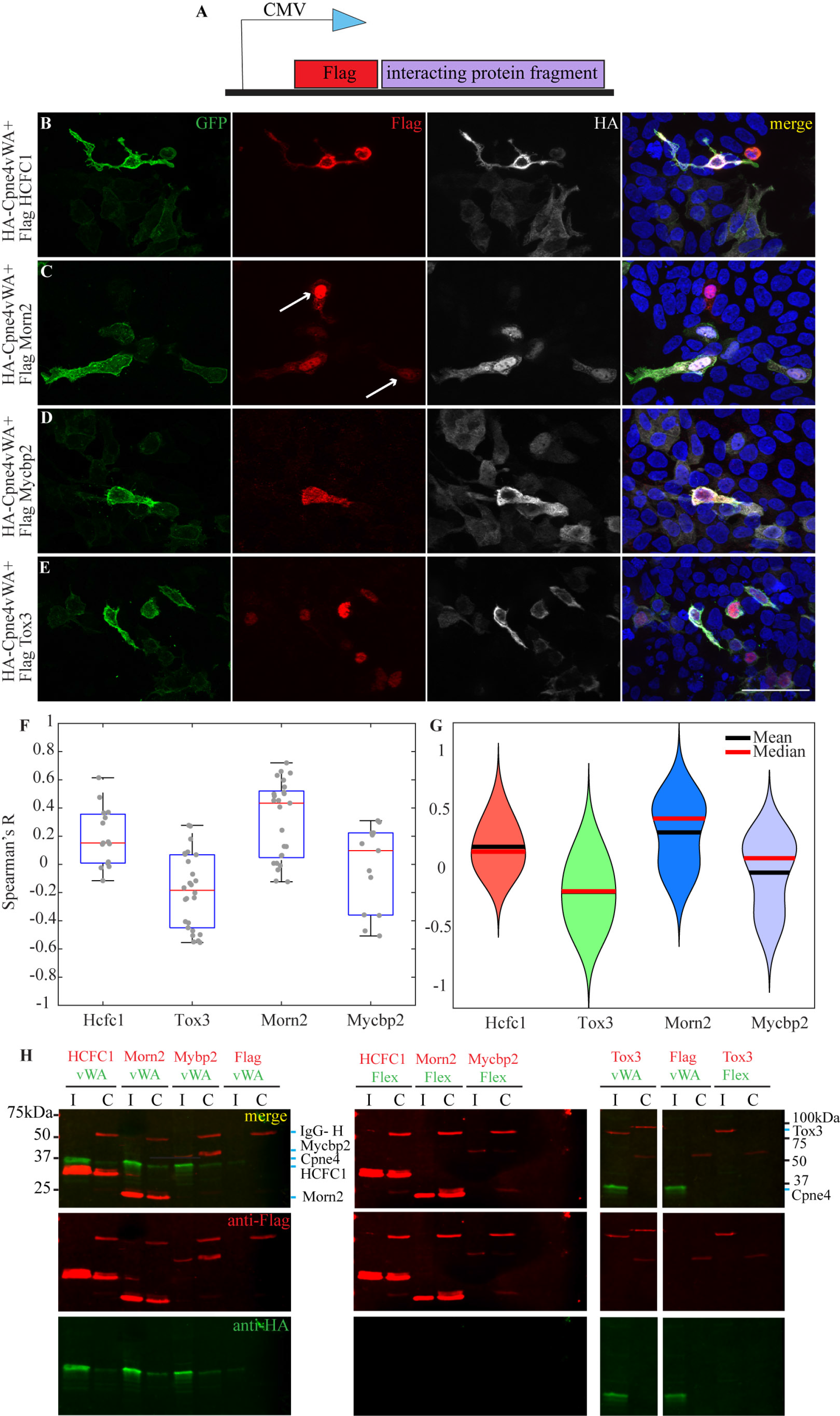
Cpne4 Adomain was pulled down with HCFC1, Morn2 and Mycbp2, but not by Tox3: (A) Map of a Flag clone consisting of a Flag tag followed by the interacting regions of HCFC1, Morn2, Mycbp2 and Tox3. (B-E) Representative images of HEK293 cells co-infected with the expression construct for Cpne4vWAdomain (HA-Cpne4vWA) and Flag-tagged interacting protein. (F, G) Box plots and violin plots indicate the Spearmann’s correlation coefficient for cololcalization between Morn2, HCFC1, Tox3 or Mycbp2 with Cpne4. (H) Western blot images of pull down from co-transfected HEK293 cells using Flag antibody show the total lysate (*I*) and the co-immunoprecipitated (*C*) proteins. Cpne4-vWA (green; 31kDa) pulled down Flag-HCFC1 (23.5kDa), Morn2 (21kDa), Mycbp2 (42kDa) and not Tox3 (66kDa). Scale bar: 50um.

**Figure 5:**
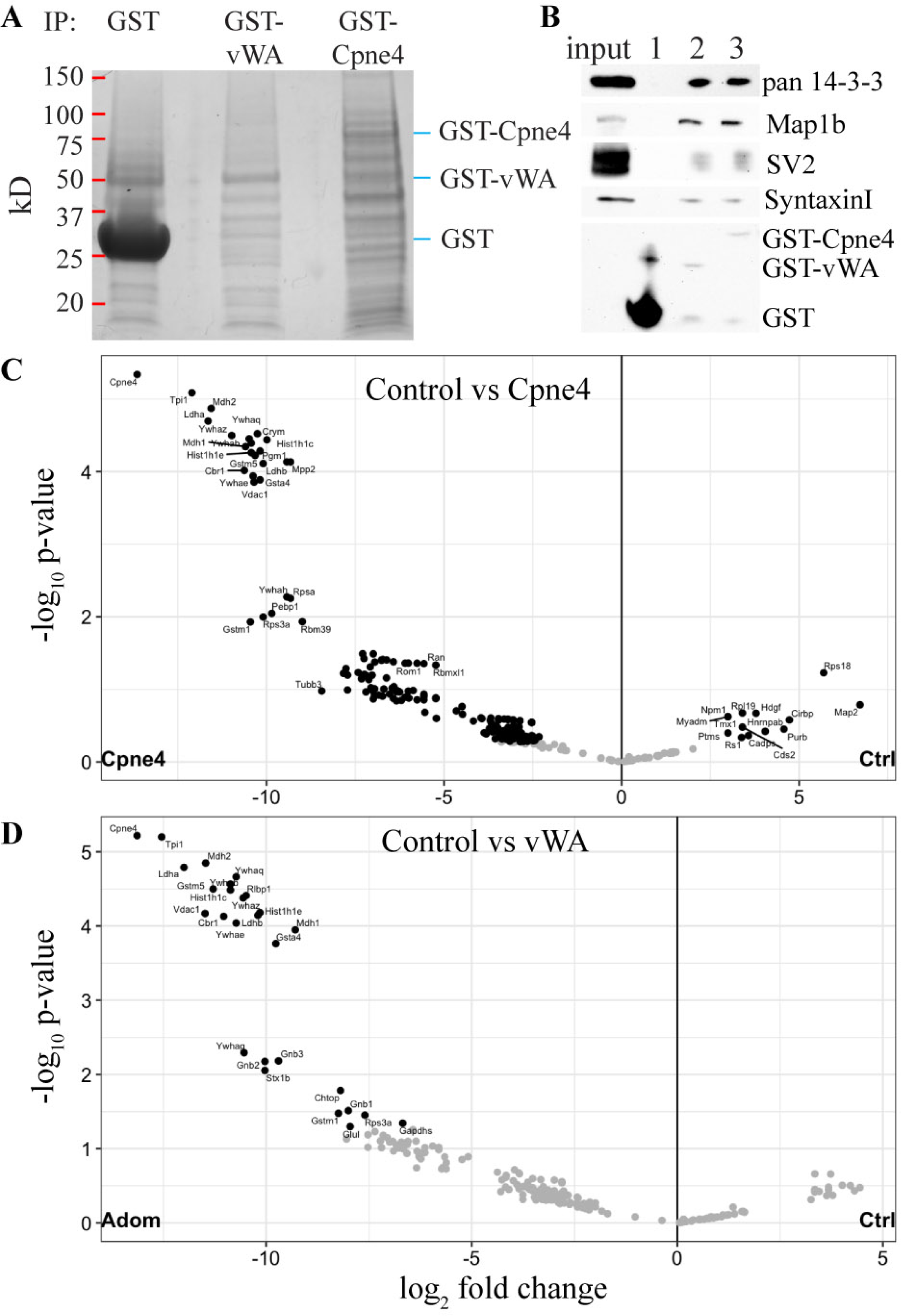
Expression of Morn2, HCFC1 and Mycbp2 in retina: (A) RNA sequencing data for Morn2, Mycbp2 and HCFC1 in different brain areas, Brn3a WT and KO RGCs, Brn3b WT and KO RGCs and rest of the retina. (B) HCFC1 (red) immunostaining in retina indicates some expression in RGCs in both the WT and Brn3b KO retinas. There is some co-labeling with Cpne4 (green) in the RGC cell bodies (arrows). (C) Morn2 (green) immunolabeling in the retina indicated mostly dendritic labeling in the IPL in both Brn3b WT and KO retinas. There is also some Morn2 labeling in the OPL of WT that is reduced in the Brn3b KO. Scale bar: 50um.

Colocalization analysis between Cpne4 and potential interactors showed that both HCFC1 and Morn2 had some degree of colocalization with Cpne4 (mean Spearmann’s correlation coefficient R= 0.19 for HCFC1, n=15 ROIs; mean R= 0.32 for Morn2, n=25 ROIs) whereas Tox3 did not (mean R= -0.19, n=26 images; Fig.4F, G). Furthermore, a pull-down analysis with Flag antibody showed that the identified Morn2 and HCFC1 domains can interact with the Cpne4 vWA domain while Tox3 does not (Fig.4H, I, J; n=4 for Morn2, n=3 for Tox3). Of the four Tomsig candidates, only Mycbp2 exhibited a small degree of colocalization with Cpne4 (mean R= 0.07, n=13 images; Fig.4D). Mycbp2 was also pulled down with Cpne4 vWA domain (Fig. 4H, I; n=4). The other candidates - Bicd2, Pitpnm2 and Sptbn1 - were not pulled down with Cpne4 vWA domain (data not shown).

RNA expression as observed by RNA sequencing analysis (Fig.5A) (1) reveals that HCFC1, Morn2 and Mycbp2 are expressed in RGCs as well as the rest of the retina at E15 and at P3 (Fig.5A). Immunostaining for HCFC1 (red) in the retina showed that it was localized in the Brn3b^+^ RGCs (green) in the ganglion cell layer in both Brn3b WT as well as Brn3b KO retinas (Fig.5B; n=2). Immunostaining for Morn2 (green) showed expression in the IPL and GCL in both Brn3b^WT/WT^ and Brn3b^KO/KO^ retinas (Fig.5C; n=2).

### 3.5 Mass Spectrometry analysis of retinal proteins identifies potential Cpne4 interactions

Copine4 protein interactions with the vWA domain may be modulated by the C2 domains. In order to identify potential native, full length Copine4 protein binding partners in the retina, we performed pulldown assays of mouse retina protein extracts with GST-vWA or GST-Cpne4 fusion proteins, alongside GST-only controls using Glutathione-Sepharose beads (Fig.6A). Each experimental condition was repeated in triplicate, and Glutathione eluates were loaded onto polyacrylamide gels, separated by electrophoresis, gel bands cut out and processed for LC-MS. A total of 2119 proteins were represented in the LC-MS results by at least one peptide (Supplementary table 1). In order to increase the stringency of our screen, we considered for analysis only proteins that were represented by at least one peptide in all three replicates of at least one of the pull-down conditions (GST-Cpne4, GST-vWA or GST only; 278 total). Differential expression was performed using the Bioconductor R package DEP (see material and methods), and a fold change of 2 and FDR of 5 % were used as thresholds for differential expression. Based on these criteria, 27 proteins were enriched in both GST-Cpne4 and GST-vWA relative to GST controls, while 180 proteins were significantly enriched in GST-Cpne4 but not GST-vWA samples relative to GST controls (Fig.6A, C, D, Supp. Table 1; n= 3 experiments, 19 retinas per experiment).

**Figure 6:**
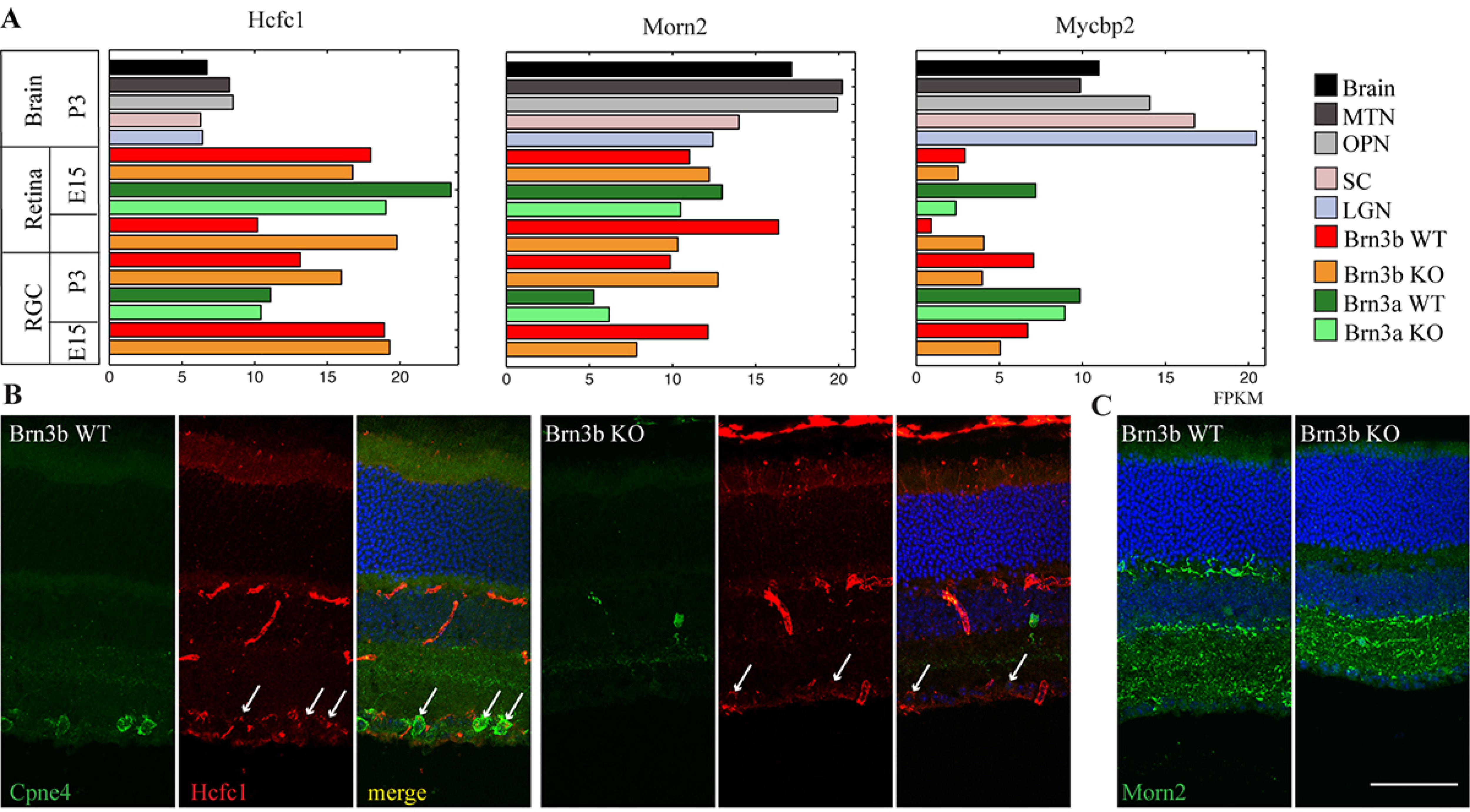
Pulldown with GST-Cpne4 or GST-CpneAdomain and LC-MS: (A) Representative SDS-PAGE gel for retina pull downs using GST-Cpne4, GST-Cpne4-vWA and GST only (n=3). (B) Representative Western blot image of some of the interactors from the pull down experiment-pan 14-3-3 (30kDa); Map1b (325kDa); SV2 (95kDa) and Syntaxin1 (33kDa) as compared to GST (28kDa) control. (C) Volcano plot for proteins that had a significant interaction with GST-Cpne4 as compared to GST control. (D) Volcano plot for proteins that had a significant interaction with GST-Cpne4-vWA as compared to GST control.

GO analysis performed on the 207 proteins that interacted with Cpne4 or Cpne4-vWA (Supp. Table 2) identified 412 biological processes and 146 molecular functions significantly enriched (FDR p<0.05). Amongst the top 30 most significantly enriched biological processes (FDR <= 5.21 e-09) all but two were related to metabolic processes, including glycolysis, ATP or nucleotide metabolism. The two biological processes unrelated to metabolism but interesting from the perspective of Cpne4 involvement in cell morphology and differentiation were “regulation of localization” (FDR <= 2.32 e-11) and “regulation of transport” (FDR <= 4.29 e-10). Molecular functions were mostly associated with nucleotide binding, protein binding, enzymatic or catalytic activity. Of the 180 GO cellular components that appeared enriched amongst Cpne4 interactors, at the very top were compartments associated with myelin sheath, cytoplasm, synapse, plasma membrane bounded cell projection, and neuron projection (FDRs <= 2.84 e-19). Interestingly, the highest fold enrichment and low FDR for Cpne4 and its interactors also suggested its localization in the late endosome lumen. A GO pathways analysis (PANTHER GO pathways) reveals that potential Cpne4 interactions participate overwhelmingly in a variety of neuronal seven transmembrane receptor pathways (Muscarinic, Metabotropic Glutamate Receptor, Opioid, Serotonine and Dopamine, etc.) at FDRs <= 8.03 e-05. PI3 Kinase and cytoskeletal regulation by Rho GTPase pathways were also significantly enriched. The complete list of all these suggested functions and locations from GO analysis are listed in Supplementary table 2.

In order to validate our MS screen, we analyzed several proteins associated with some of the enriched processes: 14-3-3 family of proteins, Map1b, SV2 and Syntaxin-1 (Supp fig. 3). Their association with either GST-vWA or GST-Cpne4 in the retina was confirmed by Western blotting of pulldowns from retinal lysates (Fig.6B). RNA sequencing data showed that the RNA for these candidates was enriched in the retina (Supp. Fig. 2). Using immunohistochemistry, pan 14-3-3 and Map1b (red) were found to colocalize in the Cpne4^+^ RGCs (green; Fig.7A, B; n=3 retinas each) in Brn3b WT and KO retina. SV2 and Syntaxin (red) were colocalized in the IPL of both Brn3b WT and KO (Fig. 7C, D; n=3 retinas each).

**Figure 7: Immunostaining.**
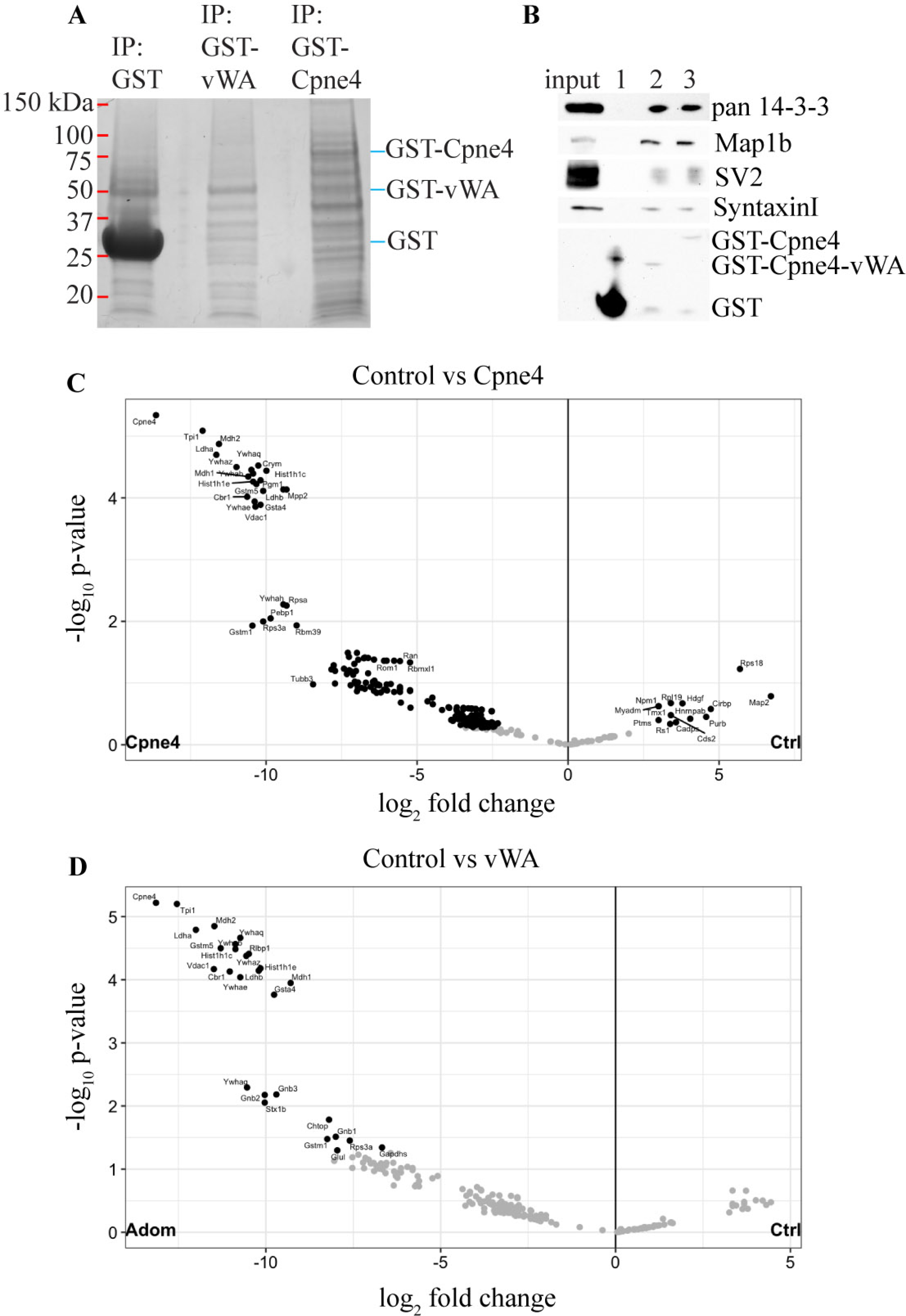
of LC-MS interactors with Cpne4 in retina: Representative images showing immunostaining of pan 14-3-3 (A); Map1b (B); SV2 (C) and Syntaxin (D; red) with Brn3b (green) in Brn3b WT (left panel) and KO (right panel). Scale bar: 50um.

## 4. DISCUSSION

Amongst Copine family members, Cpne4 has one of the highest mRNA expressions in the retina and is predominantly expressed in the RGC layer (23). Cpne4 over-expression in RGCs lead to formation of large varicosities on dendrites but no other morphological changes to RGC dendritic arbor area or stratification. The morphological defects observed by overexpression in HEK293 cells require a full length Cpne4 protein, as neither C2 domains nor vWA domain can induce extensive process formation in isolation. Cpne4 may be involved in several distinct metabolic and signaling pathways, as revealed by our Yeast two-hybrid and retina proteome interaction studies. Several proteins can interact directly with Cpne4 including Morn2, HCFC1, 14-3-3 family, Map1b, Syntaxin1 and SV2.

### 4.1 Cpne4 regulates cytoskeletal structure in the cells

It is known that Brn3 transcription factors cause morphological changes in the retina during development and these changes are most likely achieved through their transcriptional targets (2,23,27,28). Cpne4 is one such molecule that is regulated by both Brn3a and Brn3b (1). Results from the current study and previously reported experiments show that over-expression of various Copines in vitro in HEK293 cells leads to elongation of the transfected cells and formation of neurite like structures (23). Transfection with the dominant negative forms of Cpne4 however, did not result in substantial morphology changes. This suggests that the association of C2 domains and vWA domain are critical for inducing phenotypic changes in vitro. Interestingly, the GO analysis of Cpne4 and its interacting proteins suggests its localization in plasma membrane bounded cell projection, and neuron projection, among other locations indicating that it might be involved in cytoskeleton – plasma membrane interactions within the cells.

Large varicosities, similar to the ones observed on RGC dendrites following Cpne4 over-expression, are reminiscent of previously described dendrite swellings, under normal and pathogenetic conditions. In Drosophila larvae, dendrite arborizations of superficial sensory neurons are heavily remodeled during metamorphosis. Prior to pruning, the dendrites tagged for elimination experience marked calcium transients, followed by increased endocytosis and formation of varicosities, like beads on a string. The beaded dendrites are eventually resorbed prior to formation of the new arbor (29,30). Similar large varicosities are also observed on the dendrites of neurons undergoing NMDA-induced excitotoxicity (31,32).

Other Copines have been previously shown to be involved in morphological changes. Cpne6 mediates cytoskeletal changes in the hippocampus during LTP (18,19). Cpne6 interacts with Rac1 to regulate the Rac1/LimK/Cofilin signaling pathway that further regulates microtubule polymerization and hence affects the spine formation in hippocampal neurons (18). Cpne3 along with Rac1 is required for cell migration during tumor metastasis (33). Copine A, a Copine homolog in Dictyostelium, interacts with actin filaments and regulates chemotaxis and adhesion in these organisms (16). CPNA1, Copine homolog in C. elegans, is located at integrin adhesion sites in the muscle cells and acts as a linker between various myofilament proteins to maintain muscle stability (14).

Taken together, these studies suggest that Copines may mediate membrane – cytoskeletal interactions in a variety of contexts. It is not clear whether the large varicosities on RGCs indicate a role of Cpne4 in dendrite remodeling, or cytoskeletal changes during dendrite development of the retina. Expression of Cpne4 and several other Copine family members increases dramatically during the first two postnatal weeks, a period of intense RGC dendritic arbor growth, differentiation and maturation of dendritic arbor at the time of eye opening. However, the varicosities appeared irrespective of whether the Cpne4 was over-expressed at P0 when the eyes are closed and the inner retina is still developing or at P14 when the eyes are open and pruning of the dendrites happens.

### 4.2 Cpne4 and endosomal/lysosomal/autophagosomal pathways

Yeast two hybrid analysis on a retina cDNA library indicated Cpne4 interacts with Morn2. Morn2 RNA is expressed in the RGCs and protein expression is seen in the RGCs and dendritic arbor in the IPL. Morn2 consists of a single conserved domain-a MORN (Membrane Occupation and Recognition Nexus) motif. Morn2 interacts with LC3 to recruit phagosomes against various bacteria in planarian flatworms (34), and is involved in cell membrane engulfment of bacteria and eventually in the invasion of bacteria into the cells (35). Other Morn repeat containing proteins with known functions are junctophilin, retinophilin (also known as Morn4 or Undertaker), PIPK (phosphatidylinositol monophosphate kinase), and Alsin2.

We and a previous study (26) have confirmed that Mycbp2 interacts with Cpne4. Mycbp2 is a Ubiquitin ligase that contains several conserved domains that are required for a variety of functions (36). Rpm-1, the C. elegans homolog of Mycbp2, regulates AMPA receptor trafficking by regulating the degradation of DLK1 which further regulates Rab5 endosomal protein (37,38). Rab5 endosomes are required for dendritic branching (39).

Copine A, in Dictyostelium also associates with vacuoles, endolysosomal organelles and phagosomes in Ca^2+^ dependent way (40). Cpne6 translocates to clathrin coated vesicles when Ca^2+^ concentration increases in the cells (6). Previous reports suggest Cpne1 involvement in endosomal and autophagocytosis pathways. Specifically, Cpne1 together with AnnexinA1 and AnnexinA5 and is involved in calcium dependent autophagosomal degradation (41,42). Cpne1 is also involved in endoproteolysis of NFƘB (43).

Further studies need to be done to see if Cpne4 is involved in similar function in the retina. One of the top hits for probable sub-cellular location of Cpne4 based on the GO analysis of its interacting proteins is the late endosome lumen. Another possible function of Cpne4 as indicated by GO analysis suggests its role in regulation of transport in the neurons. In this context, the dendritic varicosities induced in RGCs could be the result of disrupted vesicle trafficking. We note that super-resolution imaging of varicosities reveal accumulation of membranes, besides Cpne4 itself.

### 4.3 Cpne4 and neuronal maturation and synapse formation

Copines were initially discovered in the cell bodies and dendrites of hippocampus neurons and dendrites (10,17) and have been implicated in a variety of functions in the brain. Cpne6 may be involved in long term potentiation (18,19) and suppression of spontaneous neurotransmitter release (44). Cpne1 is upregulated during neural tube closure in mouse embryo, regulates neural stem cell functions during development and it is also required for progenitor cell differentiation of neurons in the hippocampus (20,21,45). The role of Copines in the development or physiology of retina is unknown. But the interactions of Cpne4 with various proteins indicate several possible functions in the retina.

Several protein interactions with Cpne4 were observed with LC-MS analysis. Some of these that were particularly interesting and were expressed in the retina specifically in the inner retina were the 14-3-3 family, Map1b, Syntaxin1 and SV2. 14-3-3 family of proteins is one of the most abundant proteins in the central nervous system. 14-3-3Ɛ, that was found to interact with Cpne4, is previously known to bind to doublecortin (Dcx) protein, prevents its degradation which further prevents neurite formation by preventing the microtubules from invading the lamellipodia (46). 14-3-3Ɛ and 14-3-3ζ are known to regulate neurogenesis and differentiation of neuronal progenitor cells in the cortex. This is achieved by regulating the catenin/Rho GTPase/Limk1/cofilin signaling pathway to promote F-actin formation (47). 14-3-3ɣ regulates Cpne1 to enhance differentiation of hippocampal progenitor cells and neurite formation in these cells (21,48). Map1b plays an important role in axon guidance and axonal branching during the development of the nervous system (49,50). Synaptic proteins Syntaxin1 and SV2, also interacting with Cpne4, are required in neurotransmitter release at the presynaptic terminal in the retina (51,52). Interestingly, according to GO analysis of Cpne4 and its interacting proteins, one of the highest probability of localization of Cpne4 is at the synapse. This also correlates with the previous reports on localization and interaction of Cpne6 with various SNARE proteins on the presynaptic membrane as well as PSD in the postsynaptic membrane (18,44).

The yeast two hybrid analysis revealed HCFC1 as an interaction of Cpne4. HCFC1 RNA and protein is also expressed in retina and RGCs. HCFC1 was initially discovered as transcriptional co-activator that along with Oct1 forms a complex with VP16 from Herpes simplex virus and help utilize the transcriptional machinery of the host cell for replicating the virus (53). In neurons, over-expression of HCFC1 leads to neuronal maturation and also limits neurite growth (54). Mutations in HCFC1 protein are associated with Non syndromic Intellectual Disability in humans (55).

The functions of Mycbp2 in the central nervous system have been studied extensively before (36). In retina, Mycbp2 RNA expression is observed in RGCs and other cell types and is required for axonal growth cone formation (56). Knocking out Mycbp2 prevents RGC axons from reaching the thalamus and mis-targeted axons in mouse LGN (57–59). Highwire, the homolog of Mycbp2 in Drosophila is also required for synapse formation (60,61).

## 5. CONCLUSIONS

To summarize, Cpne4 over-expression leads to morphological changes in the retina. Cpne4 interacts with multiple proteins and is involved in several different pathways within the cells. This study points to several interesting directions, in relation to possible function of Cpne4 in the retina and the rest of the nervous system. Each specific interaction needs to be explored further to establish their roles in development and functioning of the retina.

**Table 1:**
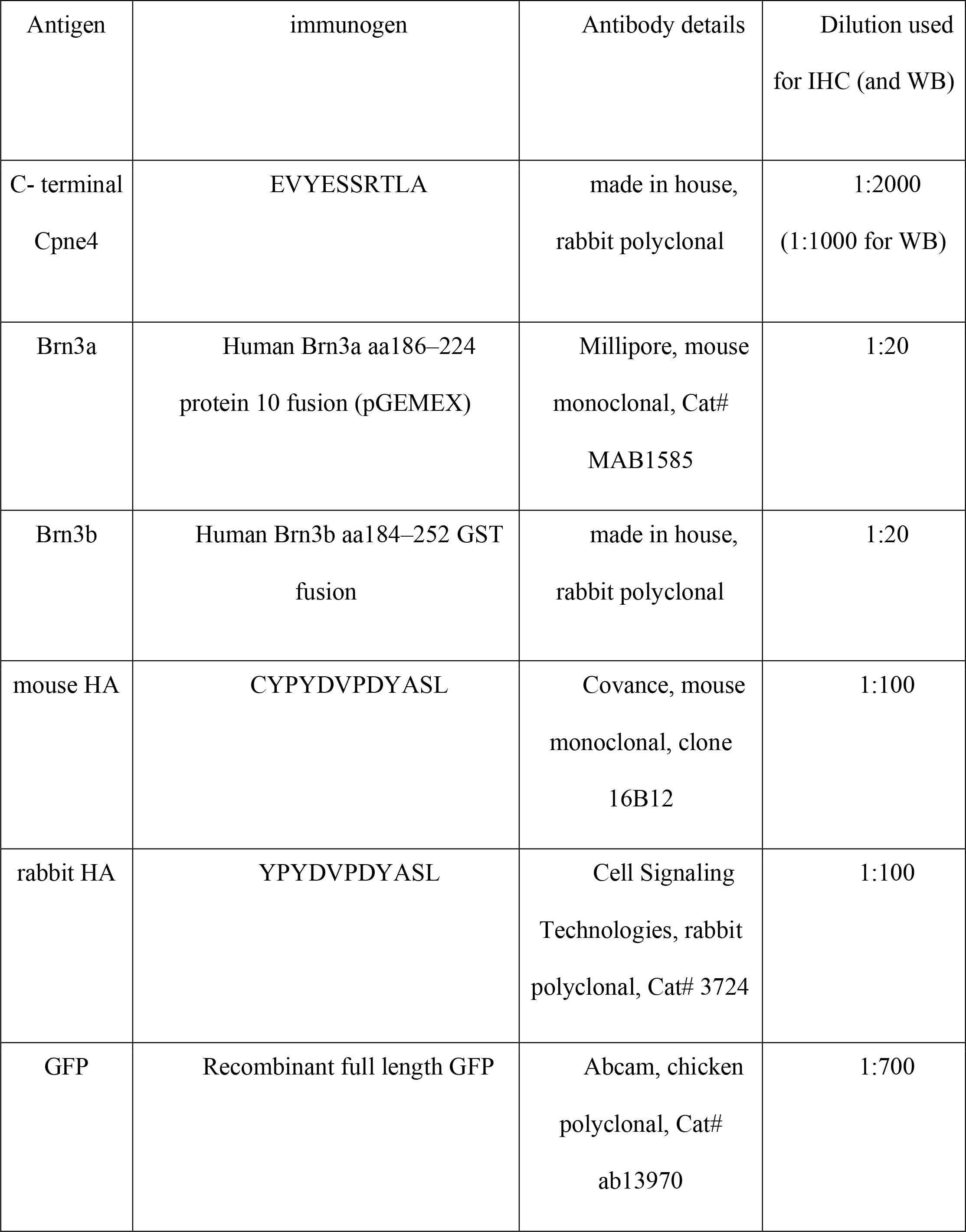
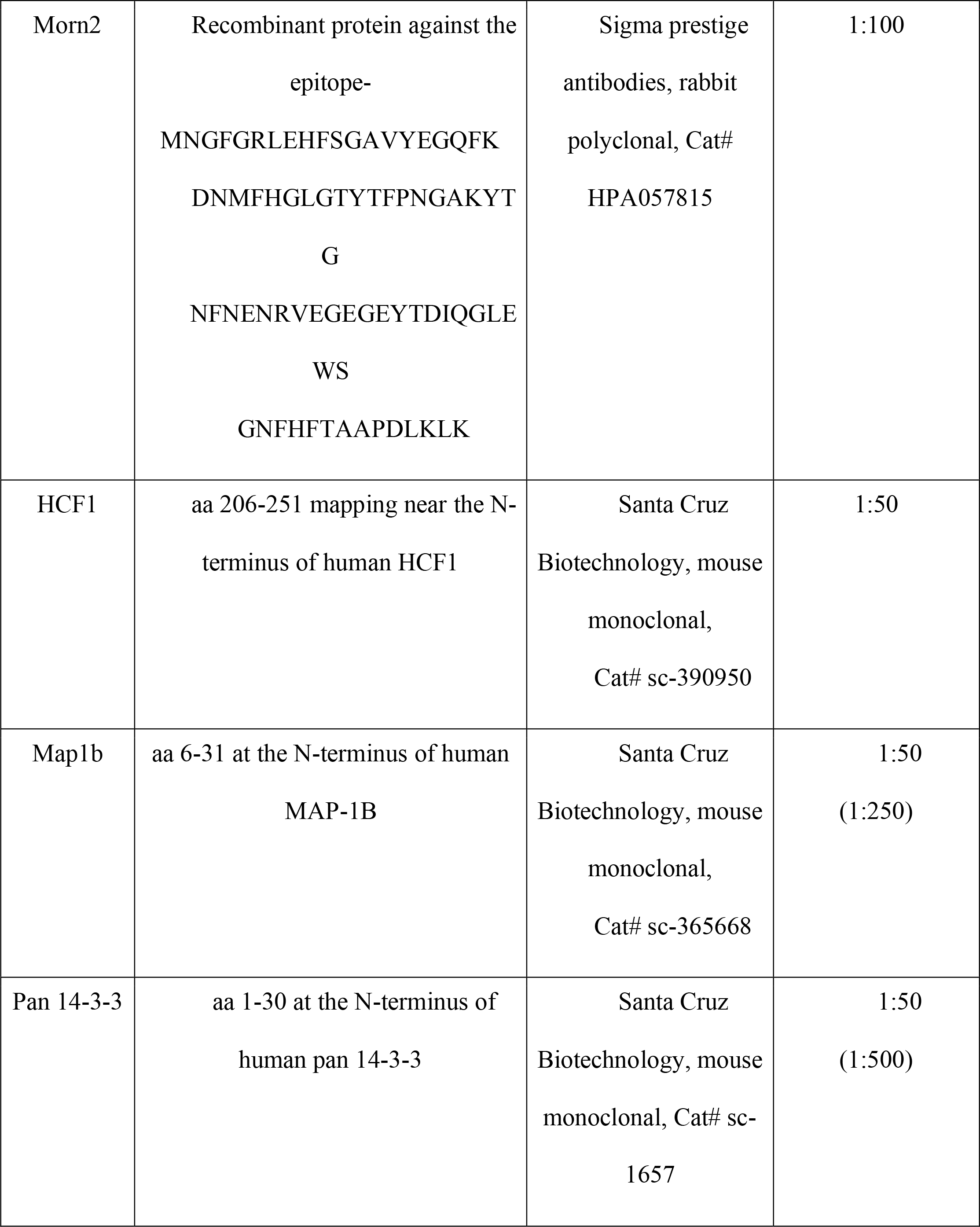
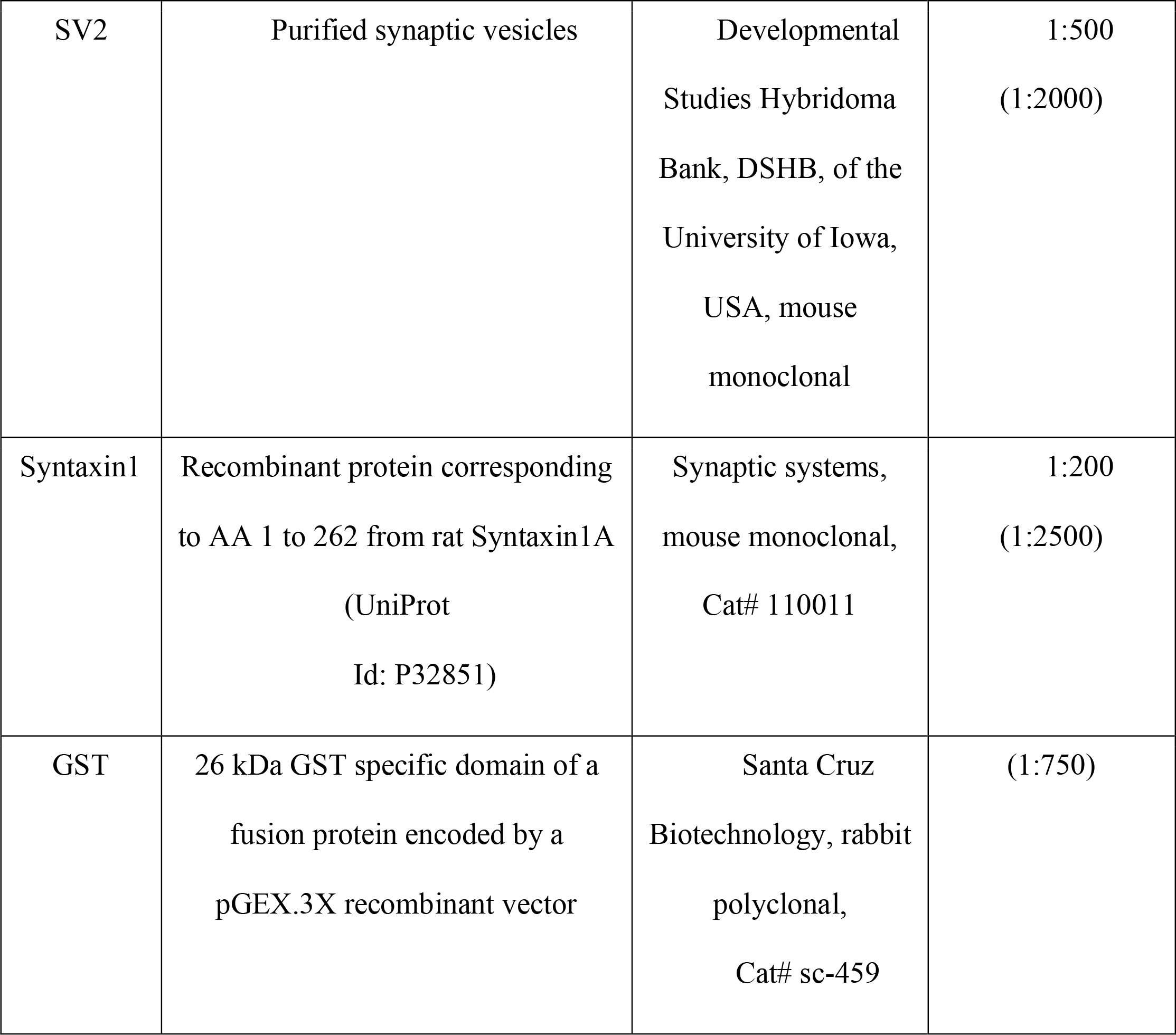
Listof antibodies used for immunofluorescence and WB

## Supporting information

SupTable1

SupTable2

## ACKNOWLEDGEMENTS

This study was funded by NIH intramural funding support to Tudor C. Badea (EY000504). We want to thank Nadia Parmhans for assistance with genotyping, Jacob Nellissery for sharing the co-immunoprecepitation protocols and Quira Zeidan (NICHD, NIH) for providing the yeast strains AH109 and Y187 and her valuable inputs on growing and mating yeast cells.

## SUPPLEMENTARY RESULTS

cDNA sequence of the interactors from the yeast two hybrid interactions-Morn2:

AAGCAGTGGTATCAACGCAGAGTGGCCATTATGGCCGGGGGAGGCCCGCTCTGTCCTAGAGCCCAGCTCCCTCCAGCCTGCCGAGCTGGCGAGCCTAGGGGAATCGCAGAGTCGCGAAAGCCTGTCCTTCACGTCTCCATCAACTTCGTCAGAAGTATATAAGATCAAATTTATATTTCCAAATGGAGACACATACGATGGCGACTGCACAAGAACTACCTCTGGGATCTGTGAGAGAAACGGGACAGGCACGCACACCACGCCGAACGGGATTGTCTACACAGGAAGCTGGAAAGATGACAAGATGAATGGCTTTGGAAGACTTGAACATTTTTCGGGCGCTGTGTATGAAGGACAGTTTAAGGACAACATGTTTCATGGACTGGGGACTTACACATTCCCAACTGGGGCAAAGTACACTGGAAATTTCAATGAAAATAGGGTAGAAGGTGAGGGAGAATACACTGACACCCAAGGCCTGCAGTGGTGTGGTAACTTCCACTTCACAGCTGCCCCCGGCCTGAAACTAAAGCTCTACATGTAGACCTGCTGCCTTAACGCTGAGATGTGGCCTCTGCAACCCCCCTTAGGCAAAGCAACTGAACCTTCTGCTAAAGTGACCTGCCCTCTTCCGTAAGTCCAATAAAGTTGTCATGCACCCACACCTTTTTGAATTATGTATTGTATGTGTGTGAGCTCATGTGTGTTTATGTGCATGTGTGTGCACATGTGCTTGTATGCATGTGTGTGTGTGTGTCTGTGTGTGAGTACTTGTTCATGCAGGTACATGTGGAAGGCAGAAGTTGATAATAAAATGCCTTTTTCAGGCTAGAACATGCTTTTAATCTCAGCACTCAGGAGAGGCAGGCAGATCTCCTGAGTTGCAGGACAGCCATGGCTACACATCTTGAAAAAGAAAAAAAAAATCCTTTTCAATCCCTTTTTACTTTTTTTTTTTTTTAAAGACATGGTCTTACTATATGGCCATGGTTGGCCTAAAACTCACTGTGTAGACCAGGTTGTCCTCAAACTCACAAAGATCTACCTGTTTCTTCTTACCAAGTGCTAGGATTAAAGGTGTTTGGTCACTAGATTCAATCTTTTCACCTCATTTGTTCCTCAGTGAACCTGGAGCTCACTGTTTCTGCTAATTTGGCTGGCCAGTGGCCCCCAGGAAACGCCCATCTCTGCTCA CCCCCAGCTGAGCACGGAGGTTAGGCACGTACCAGCGCGCCTGGAGAGGGAATCAG GGTCTTCACATTTGCAGGGAAAGCACTTTGCCCACTAAGGCATTTCCCCGACTCTTG AATTGTATCTTCATTAGGAAATGACTTTAAATAAATTGTGTGAATCAATTCAGAGTT TATGTATGTTTTAAAATTGGAGCAGTACGGTCCGAATGGTGGCTAAGTCACTATCTA ACTCTGCACCTCAACTTTTGTCAGCCCCCATGTCGGCCGCCTCGGCCTCTAGA

HCFC1: AAGCAGTGGTATCAACGCAGAGTGGCCATTATGGCCGGGGCGAGACACTCGTCGTA CCACTAACACCCCCACTGTAGTGCGGATCACTGTGGCTCCTGGGGCATTGGAGAGA GTCCAGGGTACCGTGAAGCCTCAGTGCCAAACCCAGCAGACCAACATGACCACCAC CACCATGACTGTGCAGGCCACTGGAGCTCCATGCTCAGCTGGCCCCCTGCTTAGGCC AAGTGTGGCACTGGAGTCTGGGAGCCACAGCCCTGCCTTTGTGCAACTAGCCCTTCC AAGTGTCAGAGTTGGGCTAAGTGGCCCCAGCAGCAAGGACATGCCCACAGGGCGCC AACCAGAGACATATCATACTTACACAACTAATACCCCCATGTCGGCCGCCTCGGCCT CTAGAATCCCGGCGCATGTCGGCCGCCTCGGCCTCTAGA

Tox3: AAGCAGTGGTATCAACGCAGAGTGGCCATTATGGCCGGGACCTCCAGCAAGCAAAT CAGCCACTCCCTCTCCTTCCAGCTCTATCAACGAAGAGGATGCTGATGATGCAAACA GAGCCATTGGAGAGAAAAGAACAGCCCCAGATTCTGGCAAGAAGCCCAAGACTCCA AAGAAAAAGAAAAAGAAAGATCCCAACGAGCCTCAGAAGCCAGTGTCAGCATATG CCCTGTTTTTCAGAGATACACAGGCTGCAATTAAGGGTCAAAACCCCAACGCAACCT TCGGAGAAGTCTCGAAAATCGTAGCATCTATGTGGGACAGCCTTGGGGAGGAGCAA AAGCAGGTATATAAAAGGAAAACAGAAGCTGCCAAGAAAGAATACTTGAAGGCCC TGGCTGCCTACCGGGCTAGCCTCGTTTCCAAGGCTGCTGCTGAATCTGCAGAAGCCC AGACCATCCGTTCTGTCCAGCAGACTCTGGCATCAACCAACCTGACATCCTCCCTCC TCCTGAACACATCACTGTCTCAACATGGGACAGTCCCAGCCTCACCTCAGACTCTTC CACAGTCACTCCCTAGGTCAATTGCCCCCAAACCCTTAACCATGAGACTACCCATGA GCCAGATCGTCACATCAGTCACCATTGCAGCCAACATGCCCTCGAACATTGGGGCTC CACTGATAAGTTCCATGGGGACGACCATGGTTGGTTCAGCAACCTCCACCCAGGTGA GCCCTTCGGTGCAAACCCAGCAGCATCAGATGCAGTTGCAGCAGCAACAGCAGCAG CAGCAGCAGATGCAGCAGATGCAGCAGCAGCAGTTACAGCAGCACCAAATGCATCA GCAGATCCAGCAGCAGATGCAGCAGCAGCATTTTCAGCATCACATGCAGCAGCACC TGCAGCAGCAGCATGTCGGCCGCCTCGGCCTCTAGA

## SUPPLEMENTARY TABLE TITLES

**Supplementary Table1:** Complete LC-MS data including the list of enriched proteins (Cpne4 vs control, Adomain vs control or Cpne4 vs Adomain; A) and raw data (B) containing the PSM (peptide spectrum match) values for each protein for each replicate (n=3 experiments)

**Supplementary Table2:** GO analysis of enriched proteins from LC-MS data listing most probable biological processes (A), molecular function (B), cellular component (C) and significant pathways from PANTHER analysis (D).

## SUPPLEMENTARY FIGURE LEGENDS

**Supplementary Figure 1:**
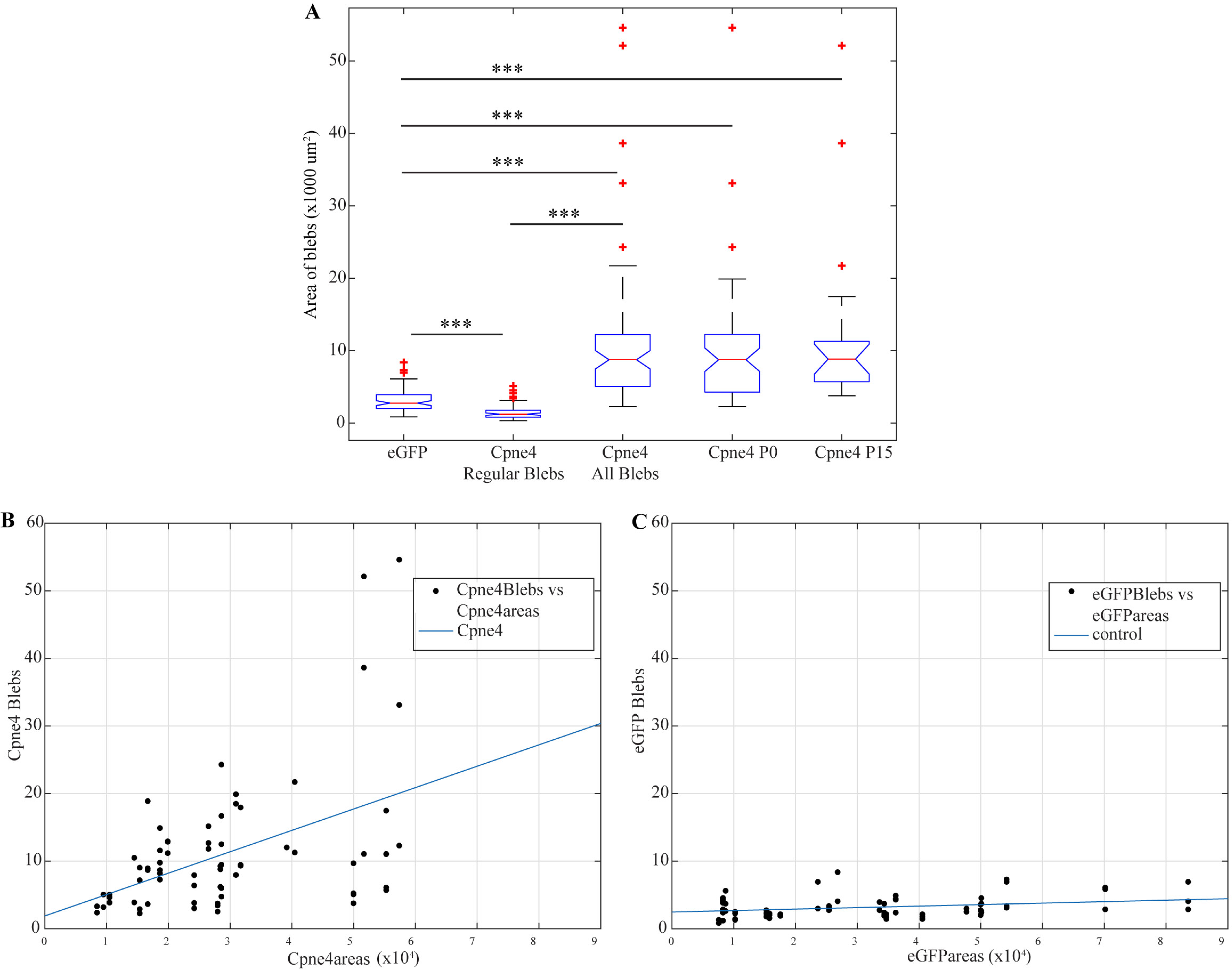
Varicosity area measurements: (A) Box plots showing the areas of the regular varicosities on eGFP infected controls, regular size varicosities on Cpne4 infected RGCs, all varicosities (regular and large blebs) on Cpne4 infected RGCs and all varicosities on P0 and P15 Cpne4 infected RGCs. (B) Plot showing correlation between arbor area and varicosity area for Cpne4 infected RGCs. (C) Plot showing correlation between arbor area and varicosity area for eGFP control infected RGCs.

**Supplementary Figure 2:**
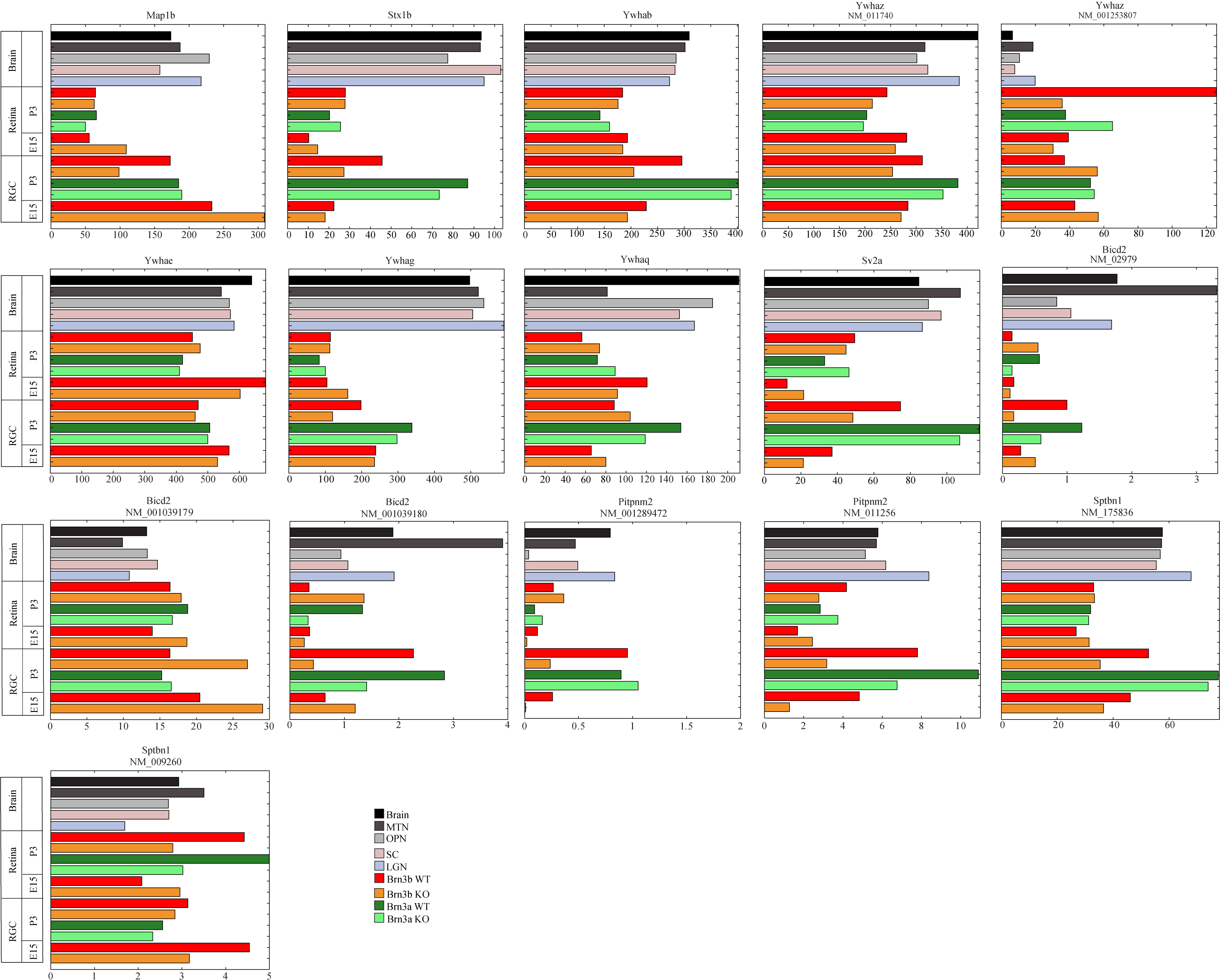
RNA sequencing data: RNA sequencing data showing Map1b, Stx1b; 14-3-3 family-Ywhab, Ywhaz, Ywhae, Ywhag, Ywhaq; SV2a; Bicd2; Pitpnm2 and Sptbn1 in different brain areas, Brn3a WT and KO RGCs, Brn3b WT and KO RGCs, rest of the retina and visual brain areas.

**Supplementary Figure 3:**
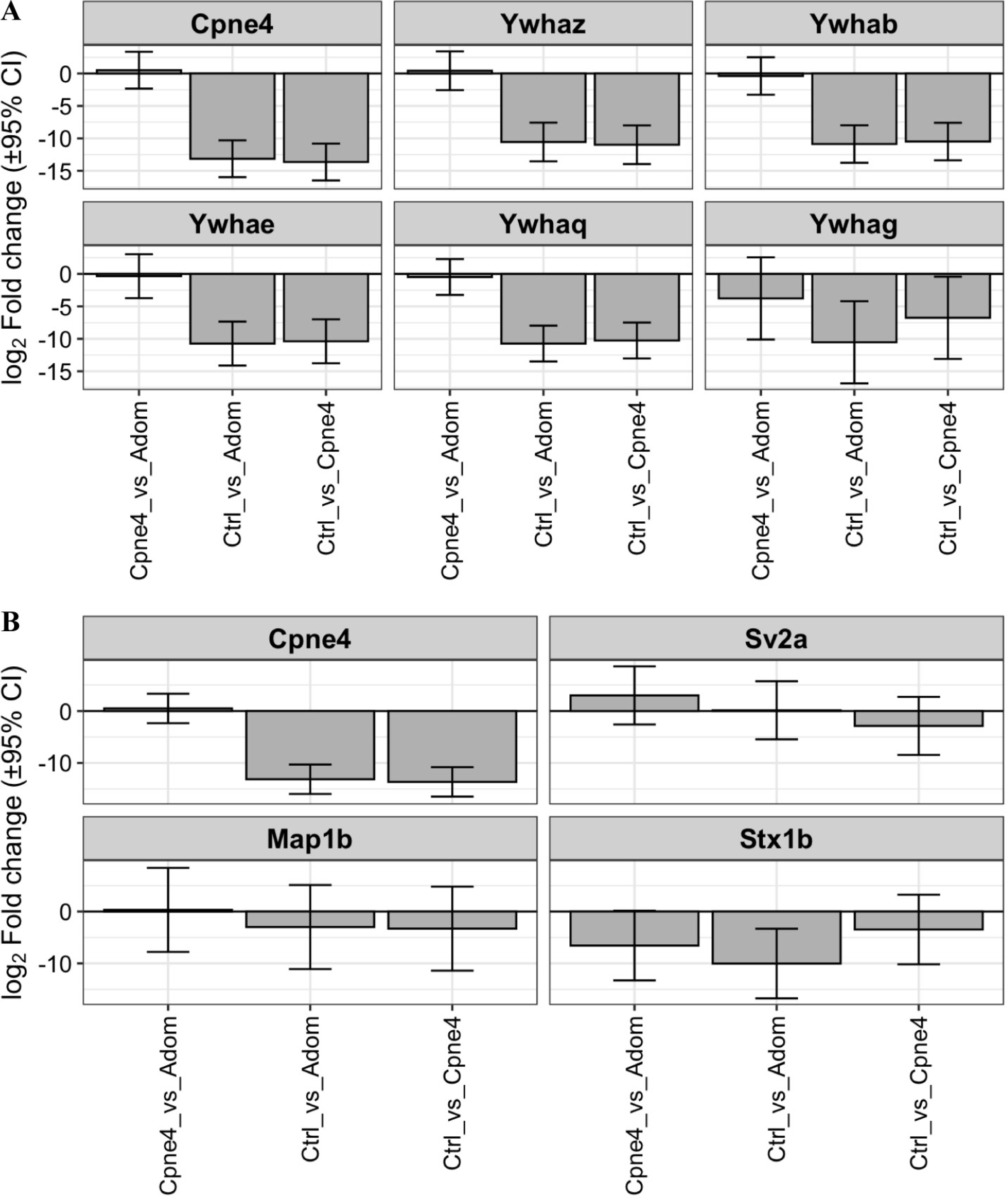
Graphs for interesting MS proteins: (A) Bar graphs showing log2 fold change as found in MS analysis for 3 different conditions-Cpne4 vs Adom (Cpne4-vWA), Ctrl (control) vs Adom and Ctrl vs Cpne4 for the proteins Cpne4, 14-3-3 family-Ywhaz, Ywhab, Ywhae, Ywhaq and Ywhag. (B) Bar graphs showing log2 fold change as found in MS analysis for 3 different conditions-Cpne4 vs Adom, Ctrl vs Adom and Ctrl vs Cpne4 for the proteins Cpne4, SV2a, Map1b and Stx1b (Syntaxin 1b).

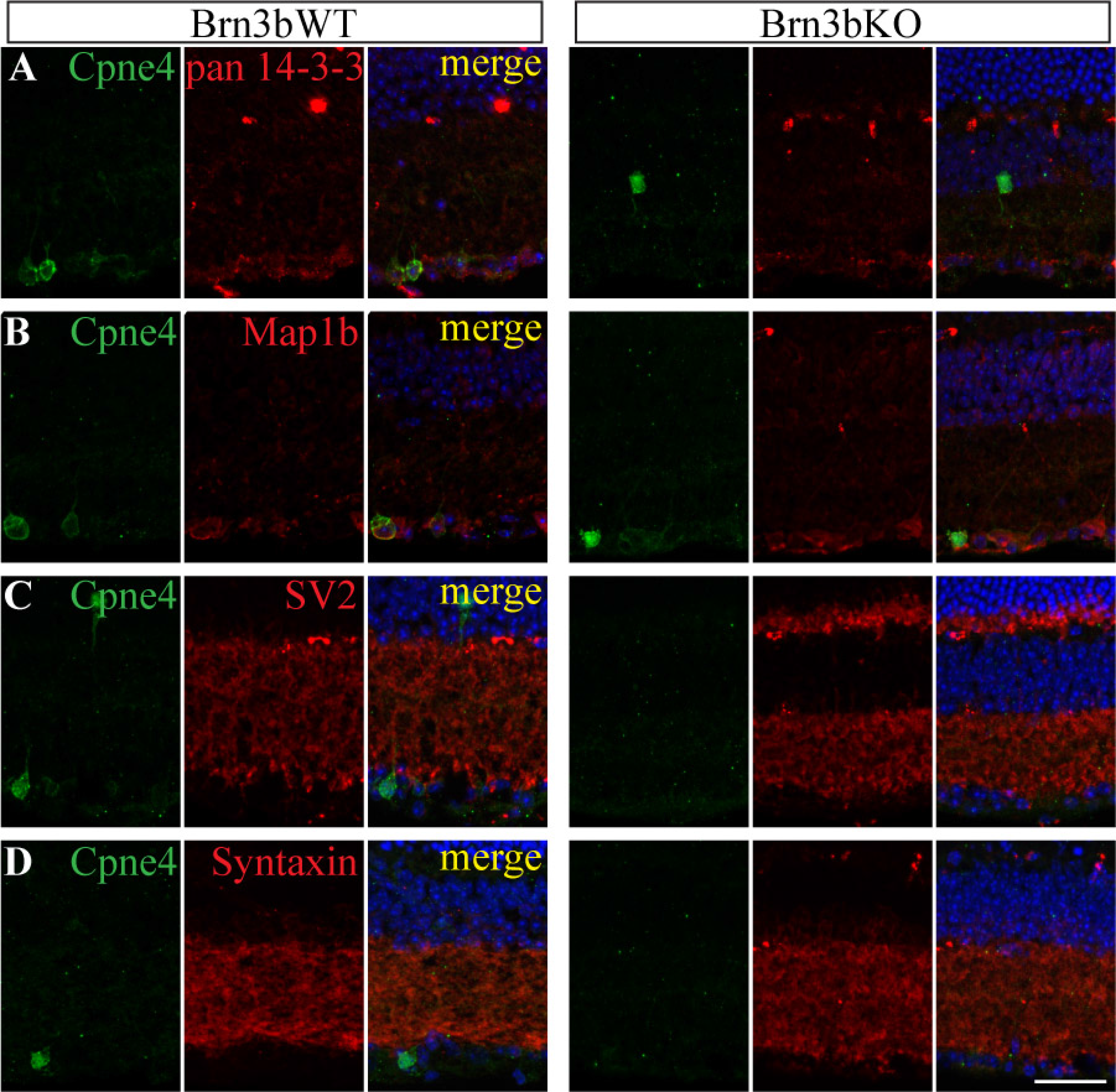

## REFERENCES

1. Sajgo S, Ghinia MG, Brooks M, Kretschmer F, Chuang K, Hiriyanna S, Wu Z, Popescu O, Badea TC. Molecular codes for cell type specification in Brn3 retinal ganglion cells. Proc Natl Acad Sci U S A. 2017;114(20):E3974–83.

2. Muzyka VV, Brooks M, Badea TC. Postnatal developmental dynamics of cell type specification genes in Brn3a/Pou4f1 Retinal Ganglion Cells. Neural Dev. 2018;13(1):1– 33.

3. Liu J, Reggiani JDS, Laboulaye MA, Pandey S, Chen B, Rubenstein JLR, Krishnaswamy A, Sanes JR. Tbr1 instructs laminar patterning of retinal ganglion cell dendrites. Nat Neurosci. 2018 May;21(5):659–70.

4. Creutz CE, Tomsig JL, Snyder SL, Gautier MC, Skouri F, Beisson J, Cohen J. The copines, a novel class of C2 domain-containing, calcium-dependent, phospholipid-binding proteins conserved from Paramecium to humans. J Biol Chem. 1998;273(3):1393–402.

5. Tomsig JL, Creutz CE. Copines: A ubiquitous family of Ca2+-dependent phospholipid-binding proteins. Cell Mol Life Sci. 2002;59(9):1467–77.

6. Perestenko P, Watanabe M, Beusnard-Bee T, Guna P, McIlhinney J. The second C2-domain of copine-2, copine-6 and copine-7 is responsible for their calcium-dependent membrane association. FEBS J. 2015;282(19):3722–36.

7. Whittaker CA, Hynes RO. Distribution and evolution of von Willebrand/integrin A domains: widely dispersed domains with roles in cell adhesion and elsewhere. Mol Biol Cell. 2002 Oct;13(10):3369–87.

8. Maitra R, Grigoryev DN, Bera TK, Pastan IH, Lee B. Cloning, molecular characterization, and expression analysis of Copine 8. Biochem Biophys Res Commun. 2003;303(3):842–7.

9. Savino M, D’Apolito M, Centra M, Van Beerendonk HM, Cleton-Jansen AM, Whitmore SA, Crawford J, Callen DF, Zelante L, Savoia A. Characterization of copine VII, a new member of the copine family, and its exclusion as a candidate in sporadic breast cancers with loss of heterozygosity at 16q24.3. Genomics. 1999;61(2):219–26.

10. Nakayama T, Yaoi T, Yasui M, Kuwajima G. N-copine: a novel two C2-domain-containing protein with neuronal activity-regulated expression. FEBS Lett. 1998 May;428(1–2):80–4.

11. Cowland JB, Carter D, Bjerregaard MD, Johnsen AH, Borregaard N, Lollike K. Tissue expression of copines and isolation of copines I and III from the cytosol of human neutrophils Abstract : Copines are a recently identified group of proteins characterized by two Ca 2 -binding C2-C terminus. Although pEST sequences indicate the. J Leukoc Biol. 2003;74(September):379–88.

12. Zhang Y, Chen K, Sloan SA, Bennett ML, Scholze AR, O’Keeffe S, Phatnani HP, Guarnieri P, Caneda C, Ruderisch N, Deng S, Liddelow SA, Zhang C, Daneman R, Maniatis T, Barres BA, Wu JQ. An RNA-sequencing transcriptome and splicing database of glia, neurons, and vascular cells of the cerebral cortex. J Neurosci. 2014 Sep;34(36):11929–47.

13. Hua J, Grisafi P, Cheng SH, Fink GR. Plant growth homeostasis is controlled by the arabidopsis BON1 and BAP1 genes. Genes Dev. 2001;15(17):2263–72.

14. Warner A, Xiong G, Qadota H, Rogalski T, Vogl AW, Moerman DG, Benian GM. CPNA-1, a copine domain protein, is located at integrin adhesion sites and is required for myofilament stability in Caenorhabditis elegans. Mol Biol Cell. 2013;24(5):601–16.

15. Ilacqua AN, Price JE, Graham BN, Buccilli MJ, McKellar DR, Damer CK. Cyclic AMP signaling in Dictyostelium promotes the translocation of the copine family of calcium-binding proteins to the plasma membrane. BMC Cell Biol. 2018;19(1):1–16.

16. Buccilli MJ, Ilacqua AN, Han M, Banas AA, Wight EM, Mao H, Perry SP, Salter TS, Loiselle DR, Haystead TAJ, Damer CK. Copine A Interacts with Actin Filaments and Plays a Role in Chemotaxis and Adhesion. Cells. 2019;8(7):758.

17. Nakayama T, Yaoi T, Kuwajima G, Yoshie O, Sakata T. Ca2+-dependent interaction of N-copine, a member of the two C2 domain protein family, with OS-9, the product of a gene frequently amplified in osteosarcoma. FEBS Lett. 1999;453(1–2):77–80.

18. Reinhard JR, Kriz A, Galic M, Angliker N, Rajalu M, Vogt KE, Ruegg MA. The calcium sensor Copine-6 regulates spine structural plasticity and learning and memory. Nat Commun. 2016 May;7:11613.

19. Burk K, Ramachandran B, Ahmed S, Hurtado-Zavala JI, Awasthi A, Benito E, Faram R, Ahmad H, Swaminathan A, McIlhinney J, Fischer A, Perestenko P, Dean C. Regulation of Dendritic Spine Morphology in Hippocampal Neurons by Copine-6. Cereb Cortex. 2018 Apr;28(4):1087–104.

20. Greene NDE, Leung K-Y, Wait R, Begum S, Dunn MJ, Copp AJ. Differential protein expression at the stage of neural tube closure in the mouse embryo. J Biol Chem. 2002 Nov;277(44):41645–51.

21. Park N, Yoo JC, Ryu J, Hong S-G, Hwang EM, Park J-Y. Copine1 enhances neuronal differentiation of the hippocampal progenitor HiB5 cells. Mol Cells. 2012 Dec;34(6):549– 54.

22. Liu C-Y, Tsai C-J, Yasugaki S, Nagata N, Morita M, Isotani A, Yanagisawa M, Hayashi Y. Copine-7 is required for REM sleep regulation following cage change or water immersion and restraint stress in mice. Neurosci Res. 2021 Apr;165:14–25.

23. Goel M, Li T, Badea TC. Differential expression and subcellular localization of Copines in mouse retina. J Comp Neurol. 2019;527(14):2245–62.

24. Gan L, Xiang M, Zhou L, Wagner DS, Klein WH, Nathans J. POU domain factor Brn-3b is required for the development of a large set of retinal ganglion cells. Proc Natl Acad Sci U S A. 1996 Apr;93(9):3920–5.

25. Zhang X, Smits AH, van Tilburg GB, Ovaa H, Huber W, Vermeulen M. Proteome-wide identification of ubiquitin interactions using UbIA-MS. Nat Protoc. 2018 Mar;13(3):530– 50.

26. Tomsig JL, Snyder SL, Creutz CE. Identification of targets for calcium signaling through the copine family of proteins. Characterization of a coiled-coil copine-binding motif. J Biol Chem. 2003;278(12):10048–54.

27. Badea TC, Cahill H, Ecker J, Hattar S, Nathans J. Distinct Roles of Transcription Factors Brn3a and Brn3b in Controlling the Development, Morphology, and Function of Retinal Ganglion Cells. Neuron [Internet]. 2009;61(6):852–64. Available from: http://dx.doi.org/10.1016/j.neuron.2009.01.020

28. Badea TC, Nathans J. Morphologies of mouse retinal ganglion cells expressing transcription factors Brn3a, Brn3b, and Brn3c: analysis of wild type and mutant cells using genetically-directed sparse labeling. Vision Res. 2011 Jan;51(2):269–79.

29. Kanamori T, Yoshino J, Yasunaga KI, Dairyo Y, Emoto K. Local endocytosis triggers dendritic thinning and pruning in Drosophila sensory neurons. Nat Commun [Internet]. 2015;6(May 2014):1–14. Available from: http://dx.doi.org/10.1038/ncomms7515

30. Williams DW, Truman JW. Remodeling dendrites during insect metamorphosis. J Neurobiol. 2005 Jul;64(1):24–33.

31. Olney JW, Fuller T, de Gubareff T. Acute dendrotoxic changes in the hippocampus of kainate treated rats. Brain Res. 1979 Oct;176(1):91–100.

32. Ahlgren H, Bas-Orth C, Freitag HE, Hellwig A, Ottersen OP, Bading H. The nuclear calcium signaling target, activating transcription factor 3 (ATF3), protects against dendrotoxicity and facilitates the recovery of synaptic transmission after an excitotoxic insult. J Biol Chem. 2014;289(14):9970–82.

33. Ahmat Amin MKB, Shimizu A, Zankov DP, Sato A, Kurita S, Ito M, Maeda T, Yoshida T, Sakaue T, Higashiyama S, Kawauchi A, Ogita H. Epithelial membrane protein 1 promotes tumor metastasis by enhancing cell migration via copine-III and Rac1. Oncogene. 2018;37(40):5416–34.

34. Abnave P, Mottola G, Gimenez G, Boucherit N, Trouplin V, Torre C, Conti F, Ben Amara A, Lepolard C, Djian B, Hamaoui D, Mettouchi A, Kumar A, Pagnotta S, Bonatti S, Lepidi H, Salvetti A, Abi-Rached L, Lemichez E, Mege JL, Ghigo E. Screening in planarians identifies MORN2 as a key component in LC3-associated phagocytosis and resistance to bacterial infection. Cell Host Microbe [Internet]. 2014;16(3):338–50. Available from: http://dx.doi.org/10.1016/j.chom.2014.08.002

35. McGuire AM, Cochrane K, Griggs AD, Haas BJ, Abeel T, Zeng Q, Nice JB, Macdonald H, Birren BW, Berger BW, Allen-Vercoe E, Earl AM. Evolution of invasion in a diverse set of Fusobacterium species. MBio. 2014;5(6):1–11.

36. Po MD, Hwang C, Zhen M. PHRs: bridging axon guidance, outgrowth and synapse development. Curr Opin Neurobiol. 2010;20(1):100–7.

37. Grill B, Bienvenut W V., Brown HM, Ackley BD, Quadroni M, Jin Y. C. elegans RPM-1 Regulates Axon Termination and Synaptogenesis through the Rab GEF GLO-4 and the Rab GTPase GLO-1. Neuron. 2007;55(4):587–601.

38. Park EC, Glodowski DR, Rongo C. The ubiquitin ligase RPM-1 and the p38 MAPK PMK-3 regulate AMPA receptor trafficking. PLoS One. 2009;4(1).

39. Satoh D, Sato D, Tsuyama T, Saito M, Ohkura H, Rolls MM, Ishikawa F, Uemura T. Spatial control of branching within dendritic arbors by dynein-dependent transport of Rab5-endosomes. Nat Cell Biol. 2008 Oct;10(10):1164–71.

40. Damer CK, Bayeva M, Hahn ES, Rivera J, Socec CI. Copine A, a calcium-dependent membrane-binding protein, transiently localizes to the plasma membrane and intracellular vacuoles in Dictyostelium. BMC Cell Biol. 2005;6:1–18.

41. Creutz CE, Edwardson JM. Organization and synergistic binding of copine I and annexin A1 on supported lipid bilayers observed by atomic force microscopy. Biochim Biophys Acta - Biomembr [Internet]. 2009;1788(9):1950–61. Available from: http://dx.doi.org/10.1016/j.bbamem.2009.06.009

42. Ghislat G, Knecht E. New Ca2+-dependent regulators of autophagosome maturation. Commun Integr Biol. 2012;5(4):308–11.

43. Ramsey CS, Yeung F, Stoddard PB, Li D, Creutz CE, Mayo MW. Copine-I represses NF-κB transcription by endoproteolysis of p65. Oncogene. 2008;27(25):3516–26.

44. Liu P, Khvotchev M, Li YC, Chanaday NL, Kavalali ET. Copine-6 Binds to SNAREs and Selectively Suppresses Spontaneous Neurotransmission. J Neurosci. 2018 Jun;38(26):5888–99.

45. Kim TH, Sung S-E, Cheal Yoo J, Park J-Y, Yi G-S, Heo JY, Lee J-R, Kim N-S, Lee DY. Copine1 regulates neural stem cell functions during brain development. Biochem Biophys Res Commun. 2018 Jan;495(1):168–73.

46. Cornell B, Wachi T, Zhukarev V, Toyo-Oka K. Regulation of neuronal morphogenesis by 14-3-3epsilon (Ywhae) via the microtubule binding protein, doublecortin. Hum Mol Genet. 2016 Oct;25(20):4405–18.

47. Toyo-oka K, Wachi T, Hunt RF, Baraban SC, Taya S, Ramshaw H, Kaibuchi K, Schwarz QP, Lopez AF, Wynshaw-Boris A. 14-3-3ε and ζ regulate neurogenesis and differentiation of neuronal progenitor cells in the developing brain. J Neurosci. 2014 Sep;34(36):12168–81.

48. Cheal Yoo J, Park N, Lee B, Nashed A, Lee Y-S, Hwan Kim T, Yong Lee D, Kim A, Mi Hwang E, Yi G-S, Park J-Y. 14-3-3γ regulates Copine1-mediated neuronal differentiation in HiB5 hippocampal progenitor cells. Exp Cell Res. 2017 Jul;356(1):85–92.

49. Meixner A, Haverkamp S, Wässle H, Führer S, Thalhammer J, Kropf N, Bittner RE, Lassmann H, Wiche G, Propst F. MAP1B is required for axon guidance and Is involved in the development of the central and peripheral nervous system. J Cell Biol. 2000 Dec;151(6):1169–78.

50. Bouquet C, Soares S, von Boxberg Y, Ravaille-Veron M, Propst F, Nothias F. Microtubule-associated protein 1B controls directionality of growth cone migration and axonal branching in regeneration of adult dorsal root ganglia neurons. J Neurosci. 2004 Aug;24(32):7204–13.

51. Sherry DM, Mitchell R, Standifer KM, du Plessis B. Distribution of plasma membrane-associated syntaxins 1 through 4 indicates distinct trafficking functions in the synaptic layers of the mouse retina. BMC Neurosci. 2006 Jul;7:54.

52. Wang MM, Janz R, Belizaire R, Frishman LJ, Sherry DM. Differential distribution and developmental expression of synaptic vesicle protein 2 isoforms in the mouse retina. J Comp Neurol. 2003 May;460(1):106–22.

53. Wysocka J, Herr W. The herpes simplex virus VP16-induced complex: The makings of a regulatory switch. Trends Biochem Sci. 2003;28(6):294–304.

54. Huang L, Jolly LA, Willis-Owen S, Gardner A, Kumar R, Douglas E, Shoubridge C, Wieczorek D, Tzschach A, Cohen M, Hackett A, Field M, Froyen G, Hu H, Haas SA, Ropers HH, Kalscheuer VM, Corbett MA, Gecz J. A noncoding, regulatory mutation implicates HCFC1 in nonsyndromic intellectual disability. Am J Hum Genet [Internet]. 2012;91(4):694–702. Available from: http://dx.doi.org/10.1016/j.ajhg.2012.08.011

55. Yu HC, Sloan JL, Scharer G, Brebner A, Quintana AM, Achilly NP, Manoli I, Coughlin CR, Geiger EA, Schneck U, Watkins D, Suormala T, Van Hove JLK, Fowler B, Baumgartner MR, Rosenblatt DS, Venditti CP, Shaikh TH. An X-linked cobalamin disorder caused by mutations in transcriptional coregulator HCFC1. Am J Hum Genet [Internet]. 2013;93(3):506–14. Available from: http://dx.doi.org/10.1016/j.ajhg.2013.07.022

56. Lewcock JW, Genoud N, Lettieri K, Pfaff SL. The Ubiquitin Ligase Phr1 Regulates Axon Outgrowth through Modulation of Microtubule Dynamics. Neuron. 2007;56(4):604–20.

57. Bloom AJ, Miller BR, Sanes JR, DiAntonio A. The requirement for Phr1 in CNS axon tract formation reveals the corticostriatal boundary as a choice point for cortical axons. Genes Dev. 2007;21(20):2593–606.

58. Culican SM, Bloom AJ, Weiner JA, DiAntonio A. Phr1 regulates retinogeniculate targeting independent of activity and ephrin-A signalling. Mol Cell Neurosci [Internet]. 2009;41(3):304–12. Available from: http://dx.doi.org/10.1016/j.mcn.2009.04.001

59. Vo BQ, Bloom AJ, Culican SM. Phr1 is required for proper retinocollicular targeting of nasal-dorsal retinal ganglion cells. Vis Neurosci. 2011;28(2):175–81.

60. Wan HI, DiAntonio A, Fetter RD, Bergstrom K, Strauss R, Goodman CS. Highwire regulates synaptic growth in Drosophila. Neuron. 2000;26(2):313–29.

61. Nakata K, Abrams B, Grill B, Goncharov A, Huang X, Chisholm AD, Jin Y. Regulation of a DLK-1 and p38 MAP kinase pathway by the ubiquitin ligase RPM-1 is required for presynaptic development. Cell. 2005;120(3):407–20.

62. Chuang K, Nguyen E, Sergeev Y, Badea TC Novel Heterotypic Rox Sites for Combinatorial Dre Recombination Strategies. G3 (Bethesda). 2015 Dec 29;6(3):559–71.

